# On the demographic history of chimpanzees and some consequences of integrating population structure in chimpanzees and other great apes

**DOI:** 10.1101/2024.06.14.599042

**Authors:** Camille Steux, Clément Couloigner, Armando Arredondo, Willy Rodríguez, Olivier Mazet, Rémi Tournebize, Lounès Chikhi

## Abstract

Reconstructing the evolutionary history of great apes is of particular importance for our understanding of the demographic history of humans. The reason for this is that modern humans and their hominin ancestors evolved in Africa and thus shared the continent with the ancestors of chimpanzees and gorillas. Common chimpanzees (*Pan troglodytes*) are our closest relatives with bonobos (*Pan paniscus*) and most of what we know about their evolutionary history comes from genetic and genomic studies. Most evolutionary studies of common chimpanzees have assumed that the four currently recognised subspecies can be modelled using simple tree models where each subspecies is panmictic and represented by one branch of the evolutionary tree. However, several studies have identified the existence of significant population structure, both within and between subspecies, with evidence of isolation-by-distance (IBD) patterns. This suggests that demographic models integrating population structure may be necessary to improve our understanding of their evolutionary history. Here we propose to use *n*-island models within each subspecies to infer a demographic history integrating population structure and changes in connectivity (*i.e.* gene flow). For each subspecies, we use SNIF (structured non-stationary inference framework), a method developed to infer a piecewise stationary *n*-island model using PSMC (pairwise sequentially Markovian coalescent) curves as summary statistics. We then propose a general model integrating the four subspecies metapopulations within a phylogenetic tree. We find that this model correctly predicts estimates of within subspecies genetic diversity and differentiation, but overestimates genetic differentiation between subspecies as a consequence of the tree structure. We argue that spatial models integrating gene flow between subspecies should improve the prediction of between subspecies differentiation and IBD patterns. We also use a simple spatially structured model for bonobos and chimpanzees (without admixture) and find that it explains signals of admixture between the two species that have been reported and could thus be spurious. This may have implications for our understanding of the evolutionary history of the *Homo* genus.

## 1 Introduction

Common chimpanzees (*Pan troglodytes*) and bonobos (*Pan paniscus*) are great apes found in western and central Africa, and they are the closest relatives to humans from which they diverged between 5 Mya [1, 2] and 7-8 Mya [3]. The current taxonomy of the genus *Pan* recognises bonobos as one unique species, geographically separated from common chimpanzees by the Congo river and from which it would have diverged between 0.9 and 2 Mya [2, 4, 5, 6]. Unlike bonobos, it is currently considered that common chimpanzees are further divided into four subspecies [2, 7]. Western chimpanzees, *P. t. verus*, occur in the most western part of the species geographic range, from Senegal on the west to Ghana on the east (see Fig. 1). The other three subspecies are separated from Western chimpanzees by the Dahomey gap, and their distribution ranges from Nigeria on the west to Tanzania on the east. From west to east, Nigeria-Cameroon chimpanzees (*P. t. ellioti*) are separated from Central chimpanzees (*P. t. troglodytes*) by the Sanaga river, and Eastern chimpanzees (*P. t. schweinfurthii*) are separated from Central chimpanzees by the Ubangi river (Fig. 1).

**Figure 1:**
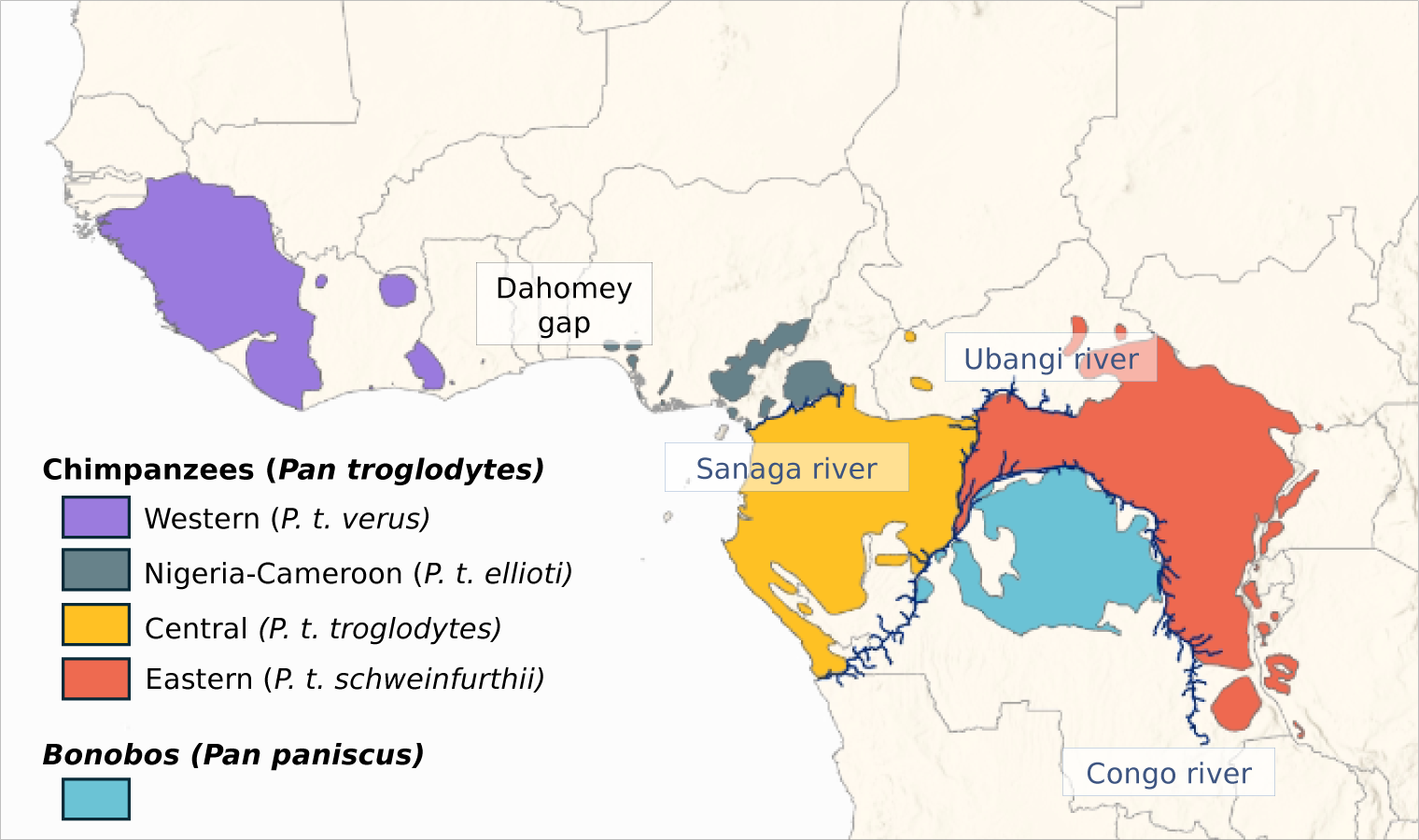
Distribution of the *Pan* genus. Data were extracted from the IUCN Red List of Threatened Species [8, 9].

Genetic and genomic analyses suggest that the four subspecies of common chimpanzees form two distinct monophyletic groups that split around 400-600 kya, with Western and Nigeria-Cameroon chimpanzees forming one clade and Central and Eastern chimpanzees forming the other clade [2, 10, 11, 12, 13, 4]. Several studies estimate that Western and Nigeria-Cameroon chimpanzees probably split between 250-500 kya [2, 6], while Eastern and Central chimpanzees separated more recently, around 90-250 kya [2, 11, 12, 6]. The existence and the magnitude of gene flow between subspecies, today or in the past, remains however unclear [13, 12, 11, 2, 14]. Indeed, if the delimitation of the subspecies reflects mostly current geographic barriers (the Dahomey gap, the Sanaga and the Ubangi rivers), it is very likely that these barriers were more or less permeable in the past [15], and it has also been suggested that the ancestral population of common chimpanzees used to cover a wider and more continuous geographic range [16]. Furthermore, studies using models of isolation with migration (IM) have found signals of gene flow between subspecies, even though there is little consensus regarding the pairs of subspecies involved (see Figure 1 in Brand *et al.* [14] for a summary). For instance, Brand *et al.* [14] have identified introgressed segments from Western chimpanzees in Eastern chimpanzees, previously suggested by Hey [11], whereas Wegmann and Excoffier [13] found gene flow from Western to Central chimpanzees. Other studies have identified isolation-by-distance (IBD) patterns, where genetic distance increases with geographic distance [17, 18]. Taking the four subspecies as a single unit, Lester *et al.* [18] computed pairwise *F_ST_* values between samples both within and between subspecies. They suggested that patterns of genetic diversity and differentiation could indeed correspond to a model with continuous gene flow across the whole *P. troglodytes* species geographic range, with the exceptions of a few highly isolated populations [18]. Patterns of IBD were also identified across Central and Eastern chimpanzees in an earlier study by Fünfstück *et al.* [17], questioning the classification of these two populations as distinct subspecies [17].

Additionally, several studies have pointed out the existence of structure within subspecies [17, 18, 19, 20, 21, 2, 6]. Central and Eastern chimpanzees could thus correspond to a set of populations that were connected by gene flow in the recent past, thus explaining the IBD pattern still detectable today [22, 17, 19]. In Nigeria-Cameroon chimpanzees, Mitchell *et al.* [20] identified two genetically distinct populations, one located in the forests of western Cameroon and another one in central Cameroon. Prado-Martinez *et al.* [2] also suggested the existence of substructure as they identified three Nigeria-Cameroon individuals who could belong to a distinct population than the rest of their sample. Finally, population structure has been identified as well in Western chimpanzees [21].

In parallel, much work has been done to reconstruct the demographic history of common chimpanzees using genetic and genomic data [2, 13, 7, 1, 11, 10, 21, 12, 23, 22, 19, 5]. Prado-Martinez *et al.* [2] used the Pairwise sequential Markovian coalescent (PSMC) method of Li and Durbin [24] on common chimpanzee individual genomes to reconstruct what Prado-Martinez *et al.* [2] interpreted as a history of effective population size (*N_e_*) changes through time. This method is particularly suited for endangered species, for which genomic data can be limited [25, 26, 27] because it only requires one single diploid genome. The interpretation of the PSMC curve is however not trivial [28, 29, 30, 31]. Indeed, whereas the interpretation of Prado-Martinez *et al.* [2] in terms of changes in *N_e_* is potentially valid, several studies have shown that ignoring population structure can lead to the inference of spurious population size changes [32, 33]. In the case of the PSMC method, Mazet *et al.* [28] have shown that under structured population models, the PSMC curve will not only be influenced by changes in *N_e_*, but also by population structure, and subsequently by changes in migration rates between populations [28, 29, 30, 31]. Given that population structure has been identified in common chimpanzees, both across the four subspecies and within subspecies, this means that there is currently no general model that would allow us to interpret the PSMC curves, while accounting for the observed patterns of IBD. Indeed, the current models of divergence represent the evolutionary history of the species and subspecies as successive splits of constant-size panmictic populations, which are incompatible with the PSMC curves. Altogether chimpanzees may represent interesting models to study ancient population structure and how it influences patterns of genomic diversity in present day populations. The current study benefits from work already done in humans but could also provide some interesting avenues of research for geneticists interested in ancient population structure in humans, by providing comparative data, and prompting similar work in other great apes.

Mazet *et al.* [28] introduced the IICR (inverse instantaneous coalescent rate), and showed that the PSMC is actually an estimate of the IICR and corresponds to changes in *N_e_* under total panmixia but not necessarily under other demographic models. Mazet and colleagues also showed in several studies that the IICR can be characterized for any model of population structure under the coalescent, which opened the way to doing demographic inference using the PSMC as a summary statistic [29, 30, 34, 31]. To that purpose, Arredondo *et al.* [31] developed a method that allows to infer the parameters of a stepwise stationary *n*-island model [35] from a PSMC curve. For example, it allows to infer the number of islands or demes, *n*, their size *N* (in diploids), and the times, *t_i_*, at which gene flow, *M_i_*= 4*Nm_i_*, may have changed by simply specifying a range of possible values for each one of these parameters. This method is implemented in SNIF (structured non-stationary inference framework).

In the present study, we ask whether it is possible to integrate population structure within each subspecies of common chimpanzees and infer a reasonable demographic history that explains the PSMC curves within one single model for each subspecies and then for the species as a whole. First, we use SNIF [31] to infer non-stationary *n*-island models for each subspecies of common chimpanzees, assuming constant deme size, using the PSMC curves generated by Prado-Martinez *et al.* [2]. At each step we validate the inference steps by generating IICR curves from the inferred demographic models and by applying SNIF to the inferred IICR curves. From the resulting inferences, we then propose a model of demographic history for the four subspecies, integrating the *n*-island models and a tree model consistent with previous research. For all inferred models (for each subspecies and for the general model), we predict genetic diversity (nucleotide diversity) and differentiation (*F_ST_*) both within and between subspecies and compare the predicted values to empirical estimates. We found that a model of structured populations with successive population splits and variable migration rates is sufficient to explain both the PSMC curves and several statistics of genomic diversity. We also find some discrepancies between the observed and predicted *F_ST_* values between subspecies and use these to identify future directions for research. In particular we suggest that models incorporating spatial structure should be explored. As a proof of concept we use a simple example of stepping-stone model and show how signals interpreted as signatures of admixture events between chimpanzees and bonobos could actually be explained by population structure alone. These results are thus of great importance for the analysis of primate genomes in general and of humans in particular, where admixture events have been inferred, and for which population structure has also been invoked as a possible explanation [36, 37, 38].

## 2 Materials and Methods

### 2.1 Data: PSMC curves

The PSMC curves of the chimpanzees used here were retrieved from the study of Prado-Martinez *et al.* [2] who kindly shared the *.psmc* files. We only kept the PSMC curves that were computed on genomes with a coverage higher than 12X. In total, this corresponded to a total of 17 individuals, namely three Eastern chimpanzees (*P. t. schweinfurthii*), four Central chimpanzees (*P. t. troglodytes*), five Nigeria-Cameroon chimpanzees (*P. t. ellioti*) and five Western chimpanzees (*P. t. verus*). The PSMC files were used to reproduce the PSMC curves (Fig. S3).

### 2.2 Inference of demographic histories for each subspecies under non-stationary *n*-island models

We used SNIF (Structured Non-stationary Inferential Framework), a freely available program (https://github.com/arredondos/snif) based on a method developed by Arredondo *et al.* [31] to infer parameters of piecewise stationary *n*-island models. SNIF assumes that the number of demes (*n*) and their size (*N* diploids) are unknown and constant through time, whereas scaled migration rates (*M* = 4*Nm*, where *m* is the proportion of migrating genes at each generation) are allowed to vary over time in a piecewise manner. Note that throughout the whole paper, deme sizes are given in number of diploid individuals.

More specifically, SNIF assumes that the PSMC can be decomposed by dividing time into periods, called “components” (*c*), during which migration rate, *M_i_* for component *c_i_*, is constant. SNIF will infer the best timing (*t_i_*) and duration (*t_i_*_+1_ *−t_i_*) for these components to fit the observed PSMC/IICR curve for a given and fixed value of *c* provided by the user. To estimate the parameters of the model, SNIF minimizes a distance computed between the observed PSMC/IICR and the IICR simulated under the piecewise stationary *n*-island model (see Arredondo *et al.* [31] for details). The user must specify *ω*, a parameter that weights the computation of the distance between observed and simulated IICR curves by giving more weight to either recent or ancient times. The size of the parameter space explored by SNIF is defined by the user, who specifies a range of values for the parameters of the model, namely the number of islands *n*, their size *N* (in number of diploids), the scaled migration rates *M_i_* = 4*Nm_i_* for each component *c_i_* with *i ∈ {*0*,…, c−* 1*}*, and the times *t_i_* (in generations) separating the components *c_i_* and *c_i_*_+1_ (for instance *t*_1_ separates the first component *c*_1_ that starts at *t*_0_ = 0 and the second component *c*_2_ that ends at *t*_2_). To scale and compare IICR curves to PSMC curves, a mutation rate (*µ*) and a generation time (*g*) are also required. The following values were used for the chimpanzee data: *µ* = 1.5 *×* 10*^−^*^8^ per bp per generation [14] and *g* = 25 years [2]. To reduce computation time and improve consistency across runs, we first ran inferences using a wide parameter space to identify the range of parameter values that were more likely to produce IICR curves reasonably close to the observed PSMC. This allowed us to then re-run the analyses on a smaller parameter space, making the optimization algorithm more efficient for the same number of iterations. For this exploratory step, the following ranges were used: *n ∈* [2; 100], *N ∈* [10; 2 *×* 10^4^], *M_i_ ∈* [0.01; 100], and *t_i_ ∈* [4 *×* 10^2^; 4 *×* 10^5^]. We also identified values for *c* and *ω* that would best describe the observed PSMC curves. We tested *c ∈ {*4, 5, 6, 7, 8*}* and *w ∈ {*0.5, 1*}* (1 being the default value). SNIF was run ten times for each combination of *c* and *ω* value, and each run used 50 iteration steps of the optimization algorithm. The inference is expected to improve as *c* increases, since it allows the algorithm to add more changes in *M* and thus better fit the observed PSMC/IICR curve. We allowed *c* to vary between four and eight because the minimum number of components required to explain two humps in a PSMC is *c* = 4 [28] and because we considered that using more than eight components might lead to over-parametrization (see next section on the validation process).

This exploratory step allowed us to significantly reduce the parameter space as can be seen in Table 1, where the ranges for *N* and *n* were halved or nearly halved. We also found that *ω*=0.5 generated the best fit to the observed data and we identified the most likely time windows (*t_i_*) for the changes of migration rates (see Table S1). The latter significantly reduced the time required by the optimization algorithm as the *t_i_* and *M_i_* values can vary over several orders of magnitude. In particular, the distance between the target and inferred IICR was on average smaller for the same number of iterations when we constrained the *t_i_* values (based on preliminary runs) than when we did not (results not shown). We finally found that using seven components for Western, Nigeria-Cameroon and Eastern chimpanzees, and eight components for Central chimpanzees provided a good balance between model complexity and increase in fit to the observed PSMC (and validation, see next section). For Western chimpanzees, we further specified not to fit the very recent past (< 20 kya), because we noticed that SNIF would try to fit the most recent increase (forward in time) in the IICR that we wanted to ignore as it differs in magnitude between the individuals of this subspecies. Table 1 summarizes the parameter space used for the analysis of all individuals of the different subspecies and for the results shown further down. Using this final parameter space, SNIF was run ten times on each PSMC curve, each run used 50 iteration steps of the optimization algorithm.

**Table 1:**
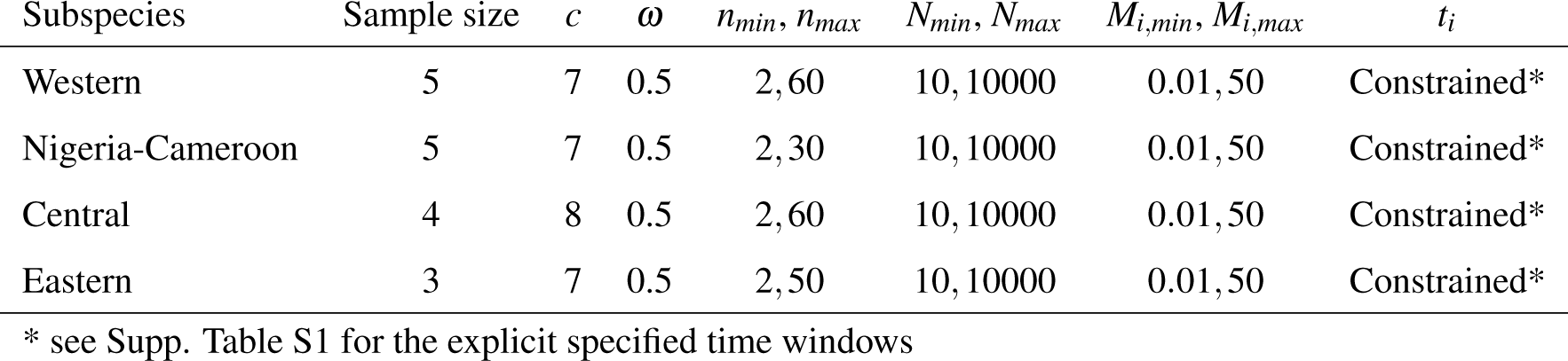
Parameter space used to run SNIF.

### 2.3 Validation of the inferred scenarios

Once we had inferred demographic scenarios, we performed a validation step as recommended by Arredondo *et al.* [31]. They suggested to simulate pseudo-observed data (POD) under an inferred scenario *S^∗^*, here in the form of IICR or PSMC curves, and to analyse these POD using SNIF with the same parameter ranges and *c* and *ω* values as those used to analyse the original observed PSMC curves. This procedure is somewhat similar to the validation process used in Approximate Bayesian Computation (ABC, [39]). In practice, SNIF allows the user to generate a *ms* command [40] to simulate coalescent times (*T*_2_) under an inferred scenario *S^∗^*. This command is used by SNIF as an input to, first, produce a pseudo-observed IICR curve using simulated *T*_2_ values, and second, infer the best stepwise stationary *n*-island model. This allows to quantify the discrepancy between the inferred model and the pseudo-observed underlying one. It is also possible to produce a *ms* command to simulate genomic data (by adapting the *ms* command using appropriate mutation and recombination rates and generation time) and run the PSMC method to produce a PSMC curve that can then be used as POD. Note that using genomic data instead of *T*_2_ values is more time consuming (several hours to produce a pseudo-observed PSMC curve) and can only be applied to a limited number of simulated scenarios. Here, we performed the validation step using both simulated IICR and simulated PSMC, as explained below and in the Supplementary material.

Different individuals of the same subspecies exhibit PSMC curves that can differ more or less significantly in the recent past (see Figure S3 and Results). In addition, running SNIF on a particular PSMC curve can generate slightly different scenarios (see Results) generally characterized by similar connectivity graphs. Instead of trying to validate many similar scenarios, we arbitrarily chose an average scenario for each subspecies based on parameter values close to the median of the distribution of the inferred values. For instance, we found that the inferred values for *n* varied between 12 and 48 for Western chimpanzees, with 50% of inferred values between 17 and 31, and we thus selected a scenario with *n* = 25 and extracted the corresponding *ms* command. This *ms* command then served to produce as many independent IICR or PSMC curves as there were individuals in that subspecies. This allowed us to quantify the variation of inferred parameter values and compare it to the variation observed when analysing the empirical data. For each run, we simulated 10^6^ *T*_2_ values to produce an IICR curve, and simulated 10 genomic sequences of 100 Mb to produce PSMC curves, adapting the *ms* command using a mutation rate of 1.5 *×* 10*^−^*^8^ [14], a recombination rate of 0.7 *×* 10*^−^*^8^ per bp per generation [4], and a generation time of 25 years [2]. We produced a (pseudo-observed) PSMC curve with PSMC [24] (flags -N 25 -t 15 -r 5 -p “4+25*2+4+6”). Altogether, these pseudo-observed IICR and PSMC curves were used by SNIF as POD, and the inference steps were done exactly as described in the previous section for the empirical PSMC curves.

### 2.4 Integration of the four subspecies *n*-island models in a general tree model

In the previous section, we inferred and validated demographic scenarios able to reproduce and fit the observed PSMC curves for each subspecies independently. Here we asked whether it was possible to integrate the four sub-subspecies into one unique demographic model that could explain the individual empirical PSMC curves, while incorporating a splitting tree based on the relationships between the subspecies as inferred by Prado-Martinez *et al.* [2] and the stepwise stationary *n*-island models within each branch of the tree, instead of assuming a panmictic population or subspecies. We constructed a scenario where an ancestral species is subdivided into *n* populations, and splits at time *T_CENW_* into two branches which are themselves subdivided in demes (Fig. 7). One of these branches will later divides into a set of demes representing the ancestor of the Central chimpanzees and another set of demes corresponding to the Eastern chimpanzees’ ancestors. The other ancestral branch becomes the ancestor to Western and Nigeria-Cameroon chimpanzees following a similar process. These splits thus generate the demes corresponding to the current four subspecies, at *T_CE_* and *T_NW_*for Central/Eastern chimpanzees and Western/Nigeria-Cameroon chimpanzees, respectively (see further, Figure 7).

SNIF cannot infer complex models involving both *n*-islands and tree models. Consequently, we built a general scenario manually, using the arbitrary “average” scenario used for the validation step above as a starting point. From the subspecies scenarios, we constructed a tree model where the subspecies *n*-island models merge (backward in time) in a way similar to that used by Rodríguez *et al.* [30]. For instance, the *n*-island models of Western and Nigeria-Cameroon chimpanzees were characterized by *n* = 25 and *n* = 13 islands respectively, and we thus merged the two sets of islands so as to use the largest *n* number for the ancestral species as suggested by Rodríguez *et al.* [30]. The same process was done for Central and Eastern chimpanzees, and then again for the ancestral branches when they merged into the most ancestral meta-population. Different values were tested for *T_CENW_*, *T_NW_* and *T_CE_* to match the times at which the PSMC curves of the sub-populations were merging backward in time, namely *T_CENW_ ∈ {*900, 800, 700*}*, *T_NW_ ∈ {*900, 800, 700*}* and *T_CE_ ∈ {*600, 500, 400*}* in kya (thousands of years ago). For each scenario we generated the IICR plots using a script developed by W. Rodriguez [29]. We used the script to simulate 10^6^ *T*_2_ values with *ms*, sampling two haploid individuals in one deme, repeating the process for each of the four current meta-populations.

### 2.5 Prediction of genomic diversity and differentiation statistics

To test whether the general treee model (Figure 7) was able to predict genetic diversity and differentiation statistics in addition to the IICR, we simulated 100 segments of 1 Mb under the four subspecies models and under variations of the general model using *ms* [40], where we allowed for the splitting times (or joining times, with time going backward) to take several values (see previous section). We used a mutation rate of 1.5 *×* 10*^−^*^8^ per bp per generation [14] and a recombination rate of 0.7 *×* 10*^−^*^8^ per bp per generation [4]. We estimated genetic diversity by computing the individual observed heterozygosity (*H_o_*) in 10 diploid individuals sampled in the present from one deme for each subspecies. We computed *H_o_* as the number of heterozygous sites divided by the total length of the simulated genomes (100*1Mb). We also computed genetic differentiation (Hudson’s *F_ST_*) between demes of the same subspecies and between demes from different subspecies, sampling ten diploid individuals per deme. Pairwise *F_ST_* were computed using original scripts from Tournebize and Chikhi [38].

Empirical values of genetic diversity were retrieved from Prado-Martinez *et al.* [2] (from Suppl. Table 12.4.1) and de Manuel *et al.* [6] (Table S4). Both studies used the same genomic data produced by Prado-Martinez *et al.* [2] and computed observed individual heterozygosity. We reported in Figure 9 their measures. Empirical values of *F_ST_* were retrieved from Fischer *et al.* [22] (computed on autosomal sequences) and Lester *et al.* [18] (computed on microsatellites). Lester *et al.* [18] only published *F^′^*, another estimate of genetic distance derived from *F_ST_*, and they kindly shared with us the original *F_ST_* values (Hudson’s estimator).

## 3 Results

### 3.1 Independent demographic history of the different subspecies

We found that by using the parameter space described in Table 1 and applying SNIF to all individual PSMCs with high enough coverage within each subspecies, we were able to produce IICR curves that were similar to the observed PSMC plots, as displayed in Figure 2 for an example and Supplementary Figures S4, S5, S6, S7 for all inferences. The inferred parameters are displayed in Figure 3. Panels A and B show the distribution of the inferred number of islands *n* and their size *N* (in number of diploid individuals), respectively (see also Table S2 and S3). The largest number of islands and the smallest deme size were inferred for Western chimpanzees, with a median *n* equal to 21 (50% of the inferred *n* values being between 17 to 31) and a median *N* equal to 305 (50% of the inferred *N* values between 239 and 335). The inferred number of islands was similar for Nigerian-Cameron and Eastern chimpanzees, with median values being 11 and 13, respectively and median *N* being 1154 and 800, respectively. Finally, for Central chimpanzees, 50% of the inferred *n* and *N* fall within 16-20 and 589-834, respectively.

**Figure 2:**
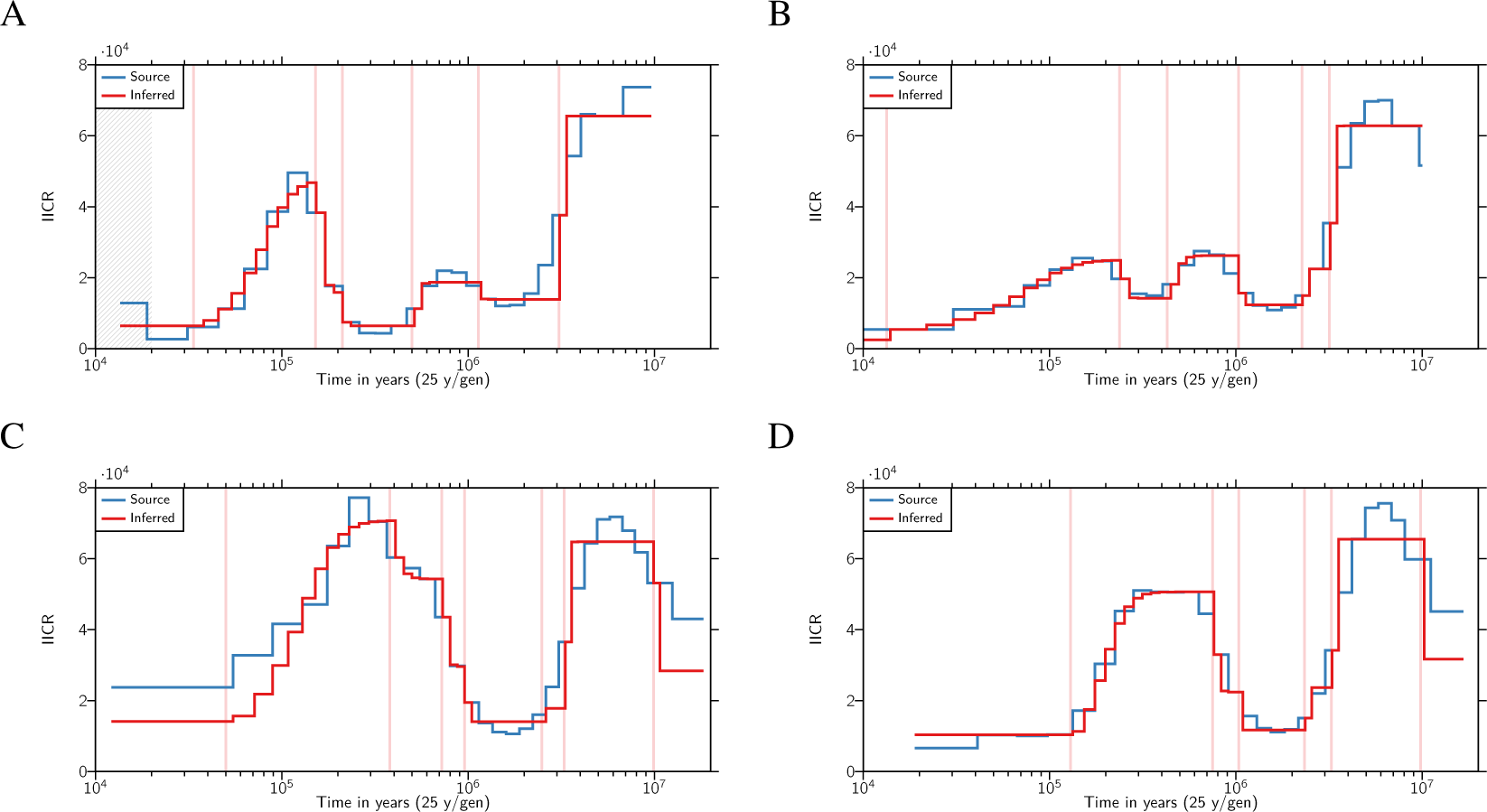
Inferred IICR curves and empirical PSMC curves for one individual per subspecies. The IICR curves are in red, and the empirical PSMC curves are in blue, and each panel corresponds to one individual from a different chimpanzee subspecies. Panel A. Western (Clint). Panel B. Nigerian-Cameroon (Damian). Panel C. Central (Vaillant). Panel D. Eastern (Kidongo). only one repetition of SNIF is displayed. See Supplementary Figures S4, S5, S6 and S7 for all the inferences. The vertical red lines highlight the times at which there is an inferred change in migration rate and therefore delimit the SNIF components. The grey zone in panel A corresponds to a part of the source PSMC which was not taken into account in the fitting of the curve by SNIF (see Material and Methods).

**Figure 3:**
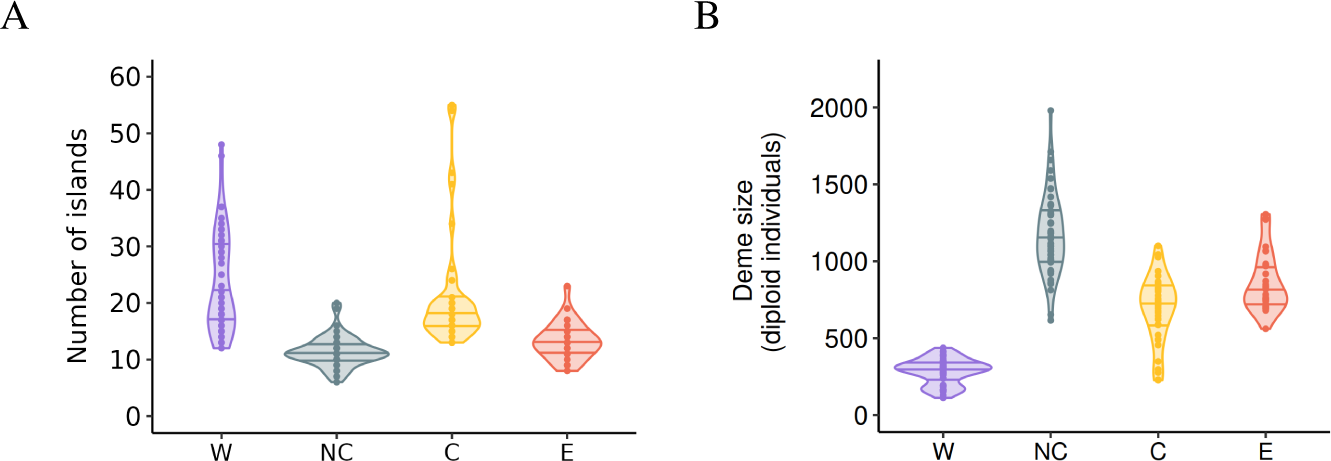
Distribution of the number and size of the demes inferred for the four chimpanzee subspecies. Panel A represents the inferred number of islands. Panel B represents the deme size (in number of diploid individuals). For both panels the results are plotted for each subspecies separately for Western (W), Nigeria-Cameroon (NC), Central (C) and Eastern (E) chimpanzees across the ten independent repetitions/inferences carried out per individual for all individuals, using the parameter space shown Table 1. Each dot corresponds to one repetition of SNIF done on one individual. The horizontal lines inside the violins correspond to the 25%, 50% (median) and 75% quantiles.

The connectivity graphs (Figure 4) show the inferred migration rates (*M_i_* = 4*Nm_i_*) through time. For the four subspecies, we observe a significant increase in connectivity (forward in time) between 2 and 3 Mya, with a higher support for the period 2.5-3 Mya, followed by a decrease around 1 Mya. This period of increased connectivity is characterized by *M_i_* values above 3 and up to 50 migrants per generation across the *n*-island for Nigeria-Cameroon, Central and Eastern chimpanzees, while values for Western chimpanzees are between 0.4 and 4. This period corresponds to a time when the four subspecies had a likely common ancestor. For Western and Nigeria-Cameroon chimpanzees, we observe a second more recent increase in connectivity between 500-600 kya and 200 kya, not observed in Central and Eastern chimpanzees, with *M_i_*values ranging between 0.8 and 50. Finally, in the more recent past, all subspecies exhibit an increase in gene flow, occurring around 100-150 kya for Eastern chimpanzees, 50 kya for Central chimpanzees, 40 kya for Western chimpanzees and between 70 and 10 kya for Nigeria-Cameroon chimpanzees.

**Figure 4:**
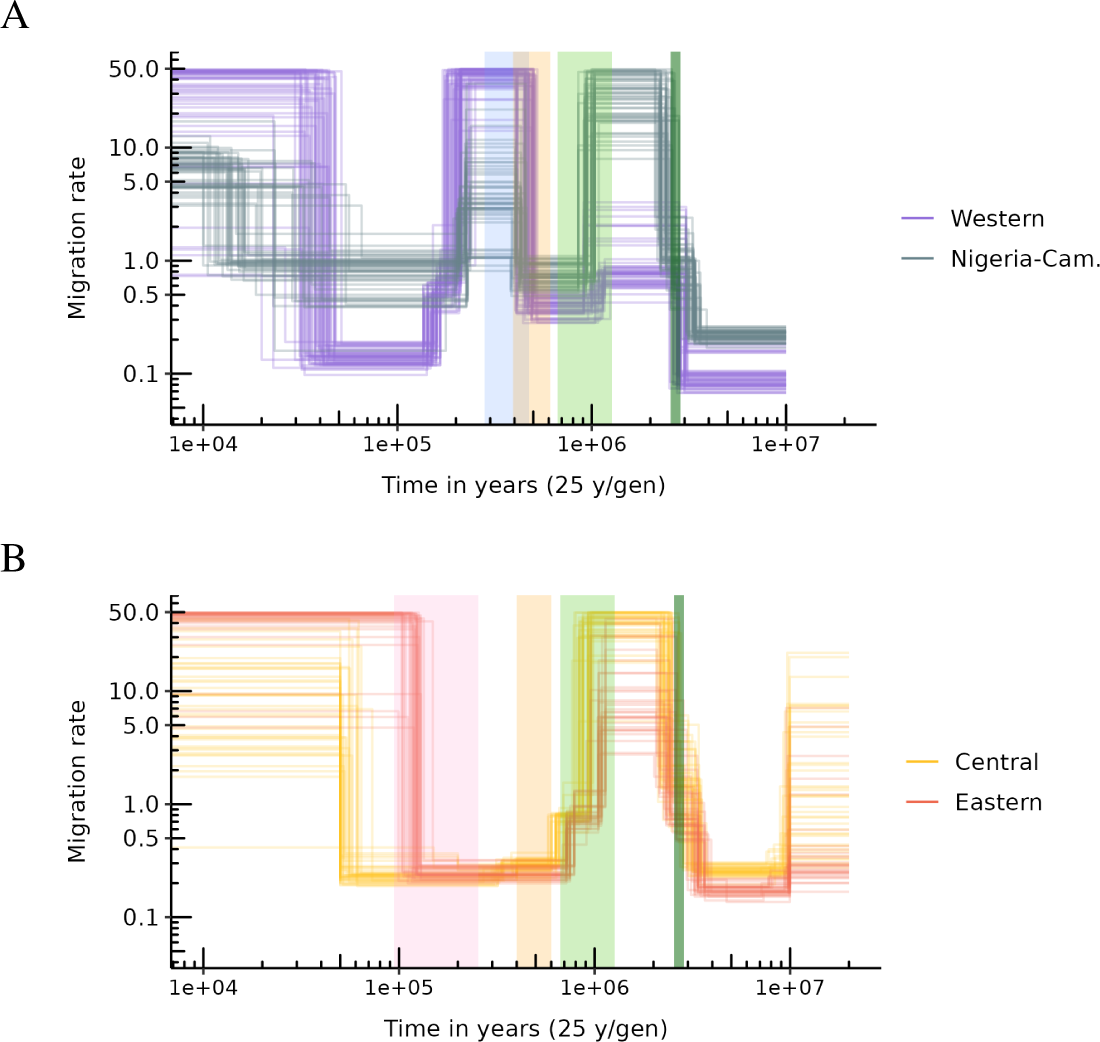
Connectivity graphs inferred by SNIF for the four chimpanzee subspecies. The y-axis represents scaled migration rates between demes (*M_i_*), and the x-axis represents time in years on a logarithmic scale. Panel A shows connectivity for Western and Nigeria-Cameroon chimpanzees. Panel B shows the same for Central and Eastern chimpanzees. Each coloured curve corresponds to one inference (one repetition of SNIF) using one PSMC curve (or individual), giving therefore 50, 50, 40 and 30 curves in total for Western, Nigeria-Cameroon, Central and Eastern chimpanzees respectively. Backward in time, the vertical coloured intervals represent respectively: C-E divergence time (in pink, panel B), W-NC divergence time (in blue, panel A), C-E and W-NC ancestral populations (in yellow, both panels), the mid-Pleistocene transition (light green, both panels) and the Pliocene-Pleistocene transition (dark green, both panels).

While the connectivity graphs are rather robust across individuals from the same subspecies and for more ancient periods for individuals from different subspecies, there is some variability in the inferred scenarios, as expected from the fact that different individuals exhibit different PSMC plots (see next section). We also observe variability in the number of islands or demes inferred for Western chimpanzees (extreme values range: 12-48 islands), and in the deme size inferred for Nigerian-Cameroon chimpanzees (extreme values range: 616-1980 diploid individuals). We also observe much variability in the most recent and most ancient parts of the connectivity graphs. For instance, there is a great variability in *M_i_* values for Central chimpanzees in the last 50 kya and before 10 Mya, and in the *M_i_*and *t_i_* values for Nigeria-Cameroon chimpanzees in the last 1 Mya.

### 3.2 Validation step

We simulated *T*_2_ values from an average scenarios identified for each subspecies and obtained IICR curves that were then analyzed using SNIF as a validation test of our inferential procedure (see Material and Methods). We were able to recover the original scenario with great precision, as shown in Figures 5 and 6. Black dots and lines are the pseudo-observed data, and the coloured patterns are the values inferred by SNIF. The inferred *n* and *N* are centered around the values that were used to generate the pseudo-observed IICRs, suggesting that SNIF is able to infer the complex scenarios inferred from the real data. We observe some variability in the inferences, as expected in any inference process and as we observed for the real data. Similarly, the inferred connectivity graphs were generally very good, with nearly no variability in the inferred *t_i_* values, and slightly more variability in the larger *M_i_* values. The shape of the inferred connectivity graphs is however perfectly inferred for the complex scenarios that had seven or eight components.

**Figure 5:**
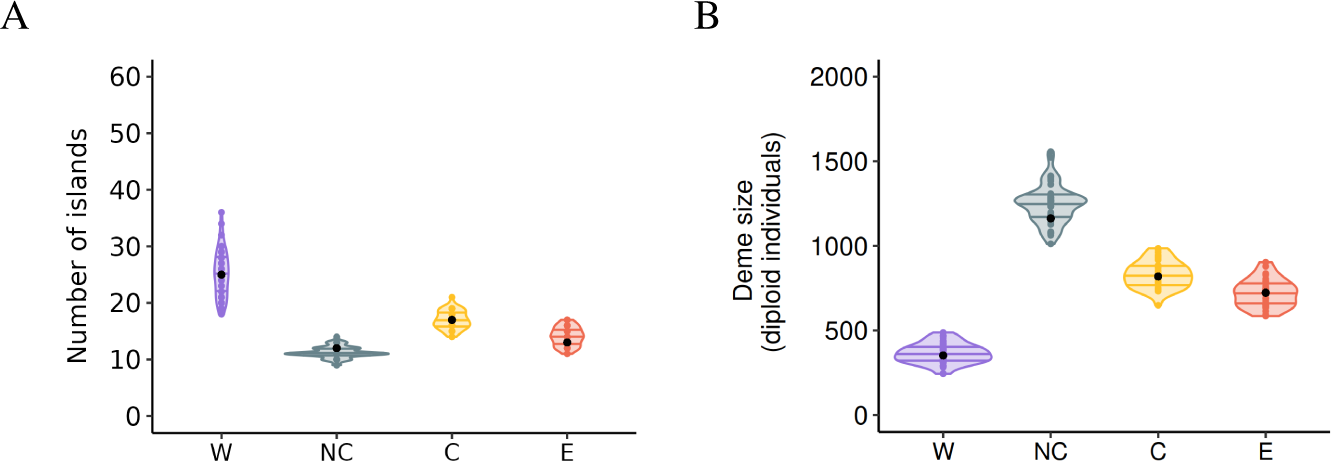
Distribution of the A. number of islands and B. deme size inferred by SNIF across the 10 repetitions per individual and all individuals for Western (W), Nigeria-Cameroon (NC), Central (C) and Eastern (E) common chimpanzees. Black dots are the pseudo-observed data.

**Figure 6:**
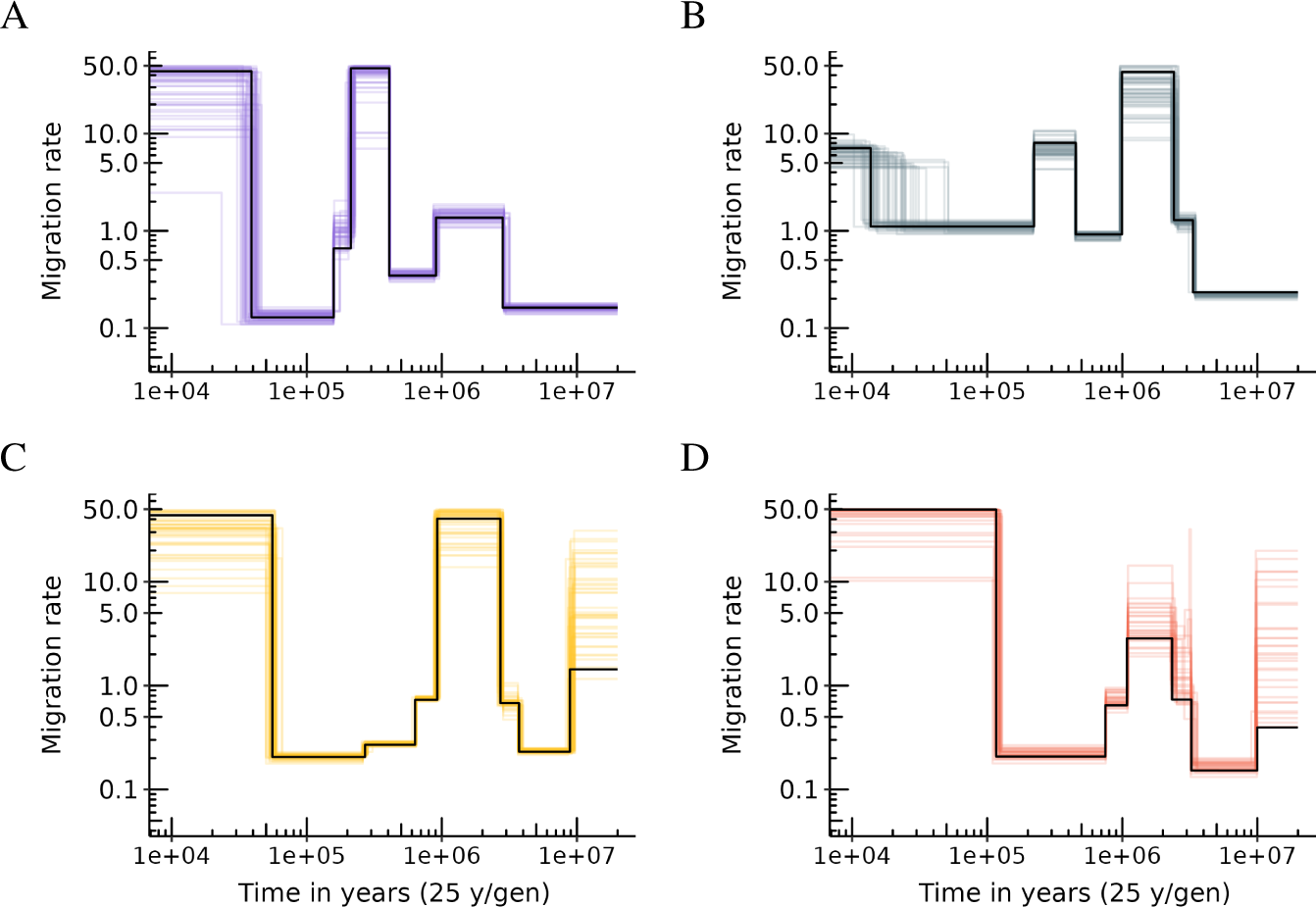
Inferred connectivity graph using a scenario inferred by SNIF as pseudo-observed data for each subspecies of common chimpanzee. A. Western chimpanzees, B. Nigeria-Cameroon chimpanzees, C. Central chimpanzees and D. Eastern chimpanzees. Black lines are the pseudo-observed data, and each colored line corresponds to one inference.

When we simulated genomic data under inferred scenarios, generated PSMC curves and provided them to SNIF as POD (Supp. Figure S10 and S11), we also recovered the original scenarios with good precision, though SNIF inferred fewer and larger islands for Eastern chimpanzees, suggesting that we might be underestimating *n* and overestimating *N* for this particular subspecies. Inferred connectivity graphs were also generally good for the four subspecies, with higher variability in the inferred *t_i_* and *M_i_*values than for pseudo-observed IICRs computed on simulated *T*_2_, which approaches the variability observed when running SNIF on empirical data. Altogether these results confirm the ability to infer a complex history of changes in connectivity under the stepwise stationary *n*-island model.

### 3.3 General *n*-island model

The general tree model incorporating within its branches *n*-island models based on the inferences presented above for each subspecies is represented Figure 7. As explained in the Materials and Methods section, it was obtained by selecting for each subspecies an “average” inferred scenario for the different subspecies. The scenarios we kept had *n* = 25 islands of size *N* = 352 (diploid) individuals for Western chimpanzees, *n* = 12 islands of size *N* = 1162 individuals for Nigeria-Cameron chimpanzees, *n* = 17 islands of size *N* = 819 individuals for Central chimpanzees and *n* = 13 islands of size *N* = 723 individuals for Eastern chimpanzees. The sets of demes were then successively divided at times corresponding to the estimated split times for the pairs of subspecies. These split times are not known but can be approximated by using the times at which the different PSMC curves join. For instance, at time *T_CE_*, the 17 demes of Central chimpanzees and the 13 demes of Eastern chimpanzees are all assumed to derive (forward in time) from the demes of their ancestor. As noted in the Materials and Methods, this ancestor was assumed to have 17 demes as 17 is the largest of the two values of *n*. For simplicity, the 13 Eastern chimpanzee demes were assumed to derive from 13 demes rather than from all 17 demes of the ancestral meta-population. Similarly, at time *T_NW_*, 12 demes of Nigeria-Cameroon chimpanzees join backward in time. 12 (out of 25) demes of the ancestor they share with Western chimpanzees. Finally, at time *T_CENW_*, the 17 islands of the ancestral meta-population of Central and Eastern chimpanzees join 17 (out of 25) islands of the ancestral meta-population of Western and Nigeria-Cameroon chimpanzees. Thus, the most ancestral meta-population is represented by 25 demes.

**Figure 7:**
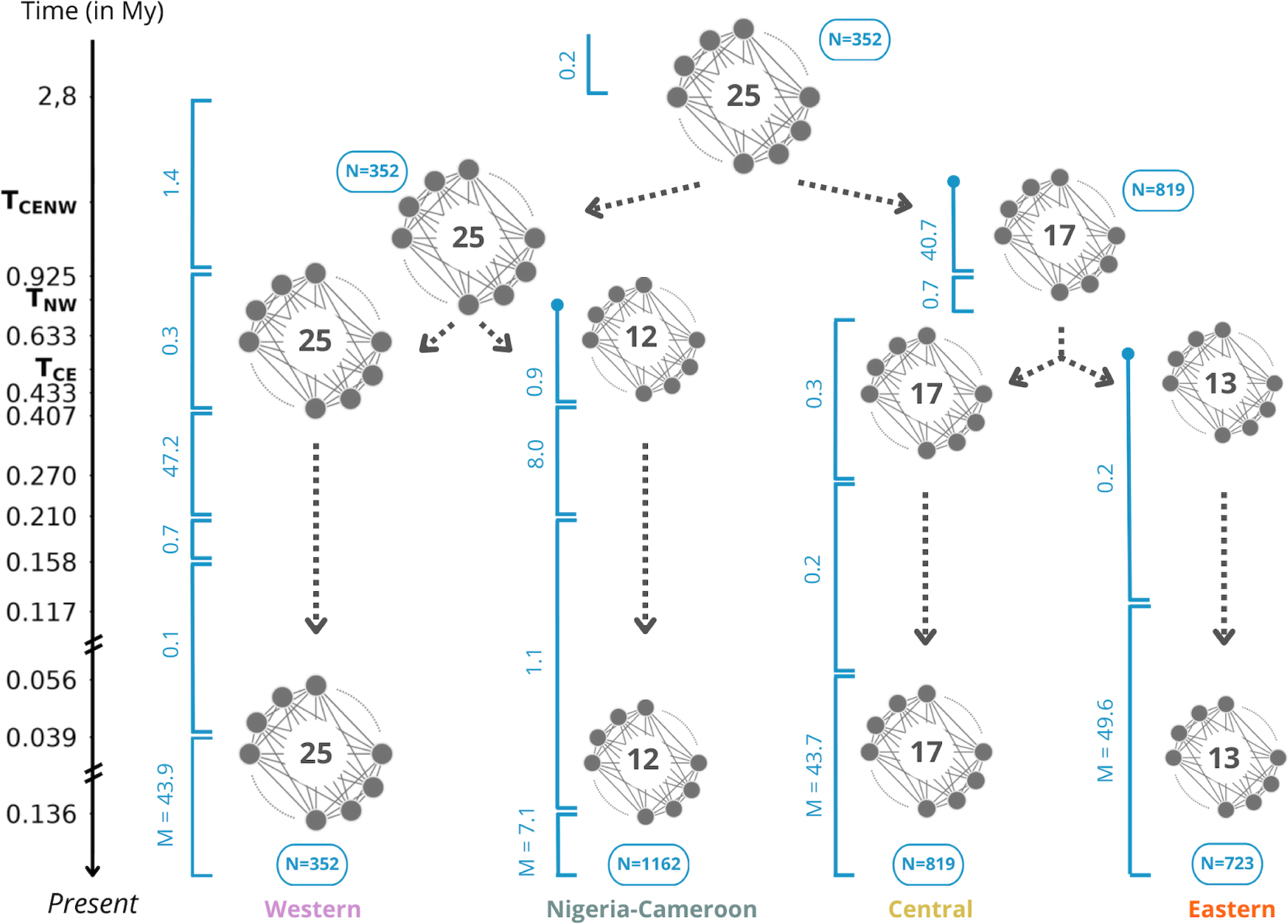
Proposed general *n*-island model for the history of the four subspecies of common chimpanzee. Dark full circles symbolize subpopulations structured in *n*-island, and the number in the middle corresponds to the number of demes *n*. Deme size *N* is given in number of diploid individuals. *T_CE_*, *T_NW_* and *T_CENW_* correspond to the times at which two meta-populations join each other (backward in time), for Central/Eastern Chimpanzees, Nigeria-Cameroon/Western chimpanzees, and the two ancestral meta-populations respectively. In blue, along the vertical intervals, are the migration rates *M_i_*= 4*Nm_i_*.

Several values for *T_CE_*, *T_NW_*, and *T_CENW_* were tested as it is unclear how closely splitting times of metapopulations correspond to splitting times of IICR or PSMC curves (see for instance Rodríguez *et al.* [30] and Chikhi *et al.* [29]). Empirical and simulated IICR curves presented in Figure 8 were obtained for *T_CE_* = 600 kya, *T_NW_* = 800 kya and *T_CENW_* = 900 kya, which are the values that gave the best visual fit of the estimated IICR curve to the observed PSMC curves (in particular for Nigeria-Cameroon and Eastern chimpanzees, see below and Supp. Figure S12-S15). For each subspecies, the simulated IICR curve produced for a sample taken in a deme is represented and closely follows the corresponding observed PSMC. We found that changing the splitting times (Supp. Figure S12-S15), and especially testing more recent values, did not significantly change the IICR curves, even though the estimated IICR could depart from the observed PSMC at the splitting time when the latter occurred too recently (see the case of Nigeria-Cameroon and Eastern chimpanzees in Supp. Figures S13 and S15 respectively). Altogether, we could construct a general model able to explain all the observed PSMC curves while incorporating both a tree model and intra-subspecies population structure, without any population size change within each branch.

**Figure 8:**
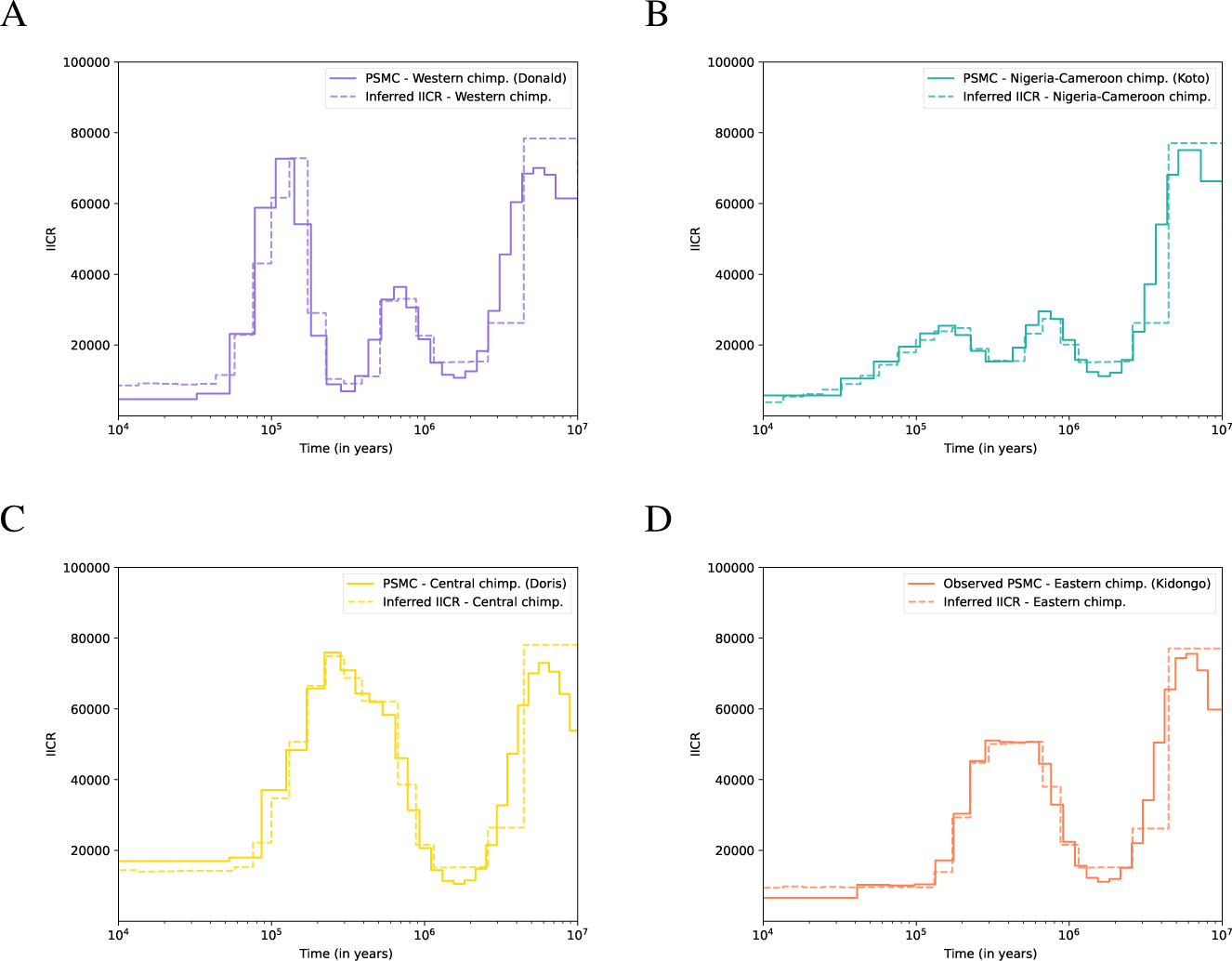
Empirical PSMC (solid line) and simulated IICR (dotted lines) for each subspecies of common chimpanzees under the n-island model displayed Figure 7, with *T_CE_* = 600 Mya, *T_NW_* = 800 Mya and *T_CENW_* = 900 Mya. A. Western chimpanzees, B. Nigeria-Cameroon chimpanzees, C. Central chimpanzees and D. Eastern chimpanzees.

### 3.4 Prediction of other genomic statistics of diversity and differentiation

Though we stress that none of the models above (individual and global) should be taken at face value, the results obtained suggest that we were able to generate IICR plots that were similar to the observed PSMC plots under 1) *n*-island models for each subspecies independently and 2) a general model incorporating the four subspecies. We simulated genomic data under our general model and computed statistics representing genetic diversity and genetic differentiation to see if it could predict values close to empirical ones. As can be seen in Figure 9, we found that our simulated diversity measures were close to the observed values computed by de Manuel *et al.* [6], and slightly lower than those computed by Prado-Martinez *et al.* [2] for Nigeria-Cameroon, Central and Eastern chimpanzees. We also found that our simulations recovered the ranking of genetic diversity for three subspecies, with Central chimpanzees being the most genetically diverse and Western chimpanzees harbouring the lowest level of genetic diversity, as observed empirically (Figure 9). However, the simulated Nigeria-Cameroon chimpanzees showed the same genetic diversity as Western chimpanzees, which is not consistent with what is found empirically [2]. These diversity estimates were identical across the different splitting times we tested.

**Figure 9:**
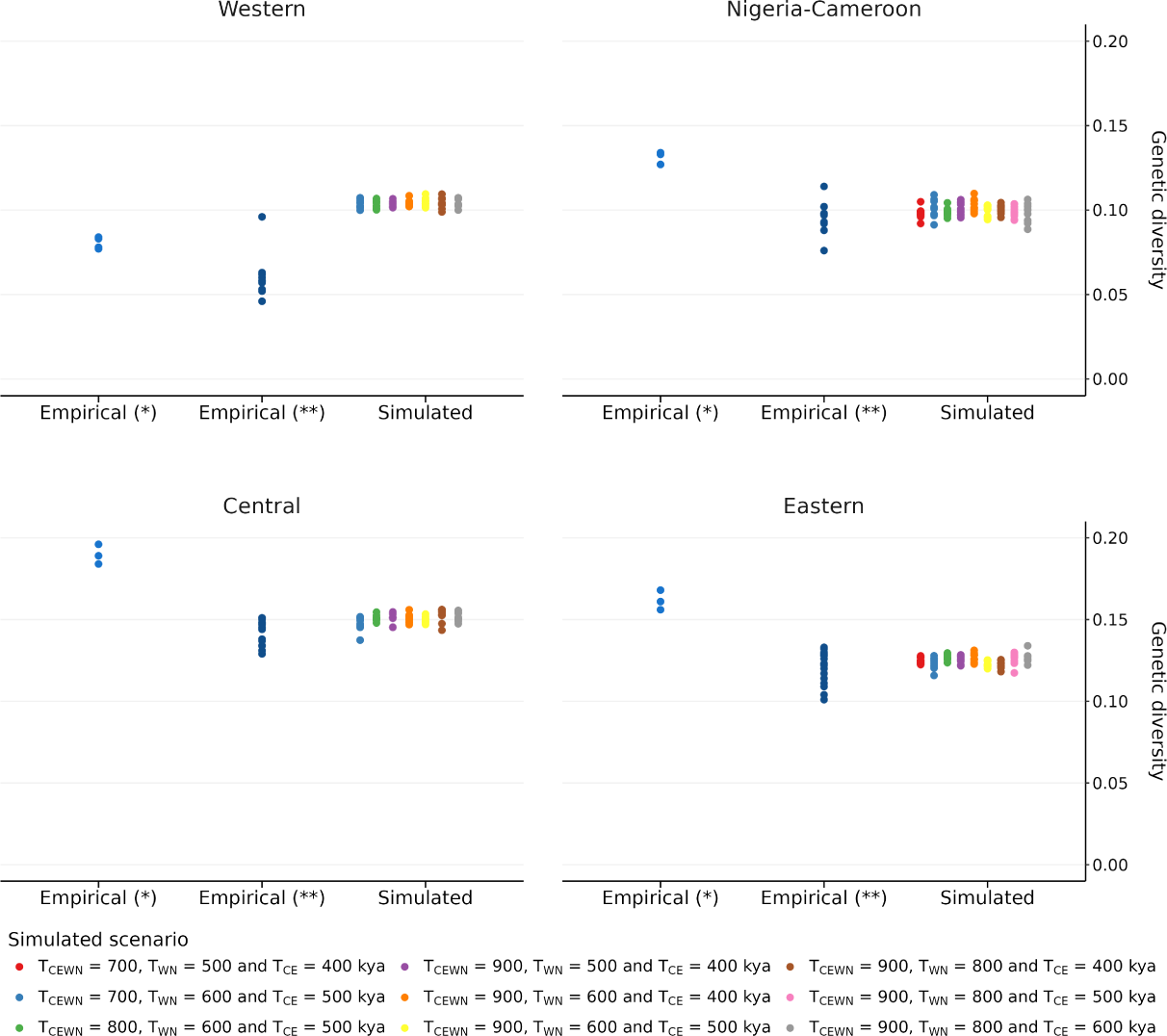
Genetic diversity of genomic data simulated under the model Figure 7 with several values for splitting times, sampling 10 diploid individuals in one deme. Empirical values were retrieved from (*) Prado-Martinez *et al.* [2] and (**) de Manuel *et al.* [6], each point corresponding to an estimate of individual observed heterozygosity.

Regarding the pairwise *F_ST_* values, Figure 10 (and Supplementary Figures S16, S17, S18) shows that the general model predicts levels of within subspecies genetic differentiation that are within the empirical distribution, with the exception of Nigeria-Cameroon chimpanzees, a subspecies for which there are nearly no observed pairwise values. Figure 10 also shows that we strongly over-estimate between subspecies differences. As expected, the *F_ST_* values between subspecies were sensitive to splitting times. For *T_CE_* = 600 kya, *T_NW_* = 800 kya and *T_CENW_* = 900 kya, the values of splitting times that follow most closely the empirical PSMC (see Figure 8), our model overestimated genetic differentiation in all pairs of subspecies. The *F_ST_* values estimated from our model were three to five times higher than their respective empirical values for all pairs of subspecies. For instance, the median of the empirical *F_ST_* ^W-NC^ was 0.12 whereas for the simulated *F_ST_* ^W-NC^ it was 0.50. For the other pairs we had similar results (median empirical *F_ST_* ^W-C^ = 0.10 vs simulated *F_ST_* ^W-C^ = 0.40, median empirical *F_ST_* ^W-E^ = 0.14 vs simulated *F_ST_* ^W-E^ = 0.46, median empirical *F_ST_* ^NC-C^ = 0.08 vs simulated *F_ST_* ^NC-C^ = 0.41, median empirical *F_ST_* ^NC-E^ = 0.08 vs simulated *F_ST_* ^NC-E^ = 0.47, median empirical *F_ST_* ^C-E^ = 0.08 vs simulated *F_ST_* ^C-E^ = 0.28).

**Figure 10:**
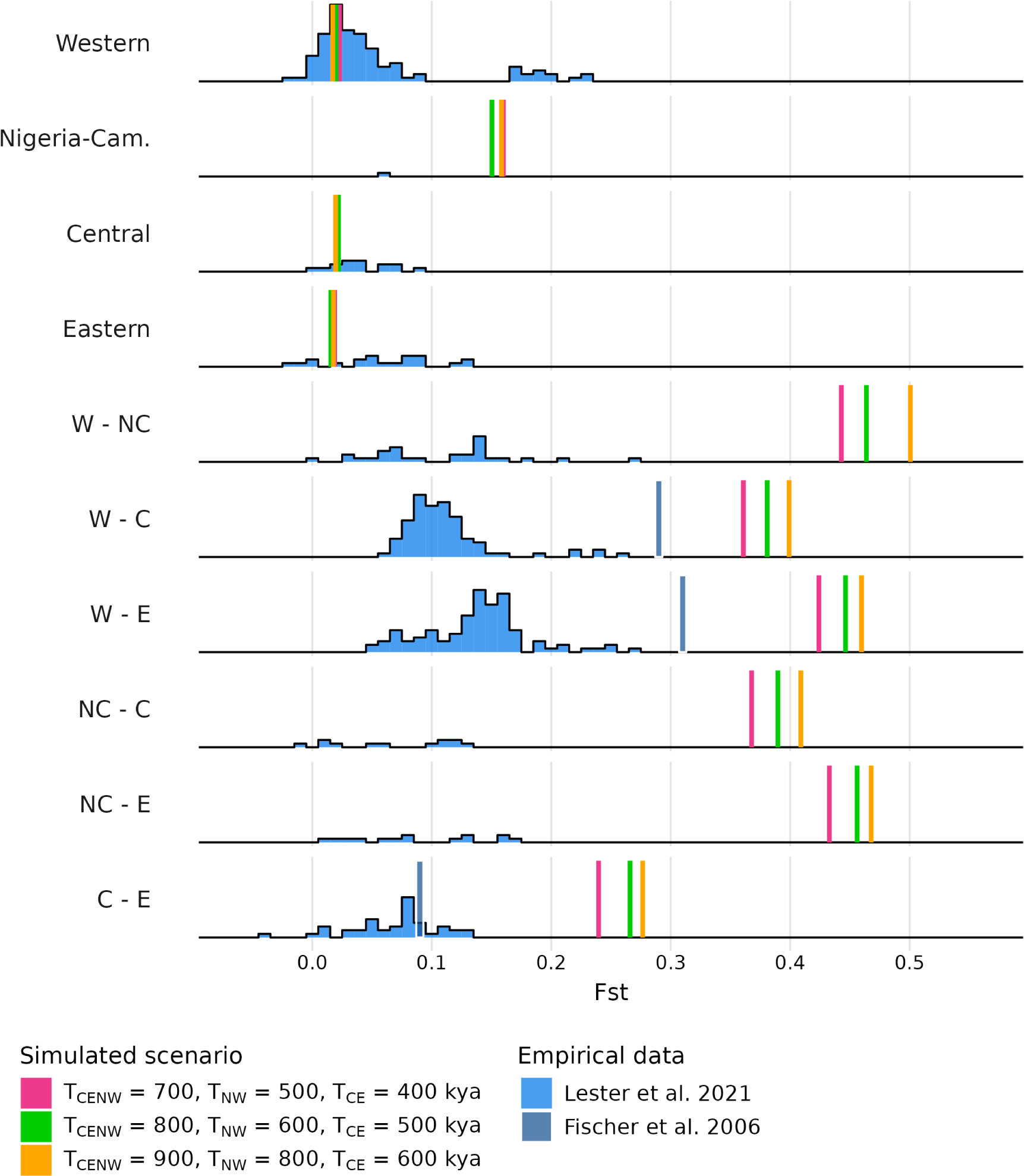
Genetic differentiation within and between subspecies. *F_ST_* were computed on genomic data simulated under the model Figure 7 with *T_CE_* = 600 kya, *T_NW_* = 500 kya and *T_CENW_ ∈ {*700, 800, 900*}* kya. See Supplementary Figures S16, S17, S18 for other splitting time values. In blue are the empirical values: histograms were retrieved from Lester *et al*. [18] (who used microsatellite data) and the blue vertical lines were retrieved from Fischer *et al*. [22] (who did not have samples of Nigeria-Cameroon chimpanzees in their study).

As expected we found that having more recent splitting times reduced our *F_ST_* estimates which were getting closer to empirical values, though they never reached them for the parameter values we tested and which were selected to match the splitting times in the PSMC curves (Supplementary Figures S16, S17, S18). Genetic differentiation between demes of the same subspecies was much lower than between demes from different subspecies and were low and similar for three subspecies (Western, Central and Eastern) but much higher value within the Nigeria-Cameroon subspecies.

As expected as well, our general model predicted lower *F_ST_*values between Western and Nigeria-Cameroon chimpanzees and between Central and Eastern chimpanzees than between other pairs of subspecies. This what is empirically observed, but we note that this is the reason why the original authors proposed the topology that we used. Finally, Eastern chimpanzees are the most differentiated subspecies to both Western and Nigeria-Cameroon chimpanzees, and Nigeria-Cameroon chimpanzees are the most distant subspecies to Central and Eastern chimpanzees, in both observed and simulated estimates.

## 4 Discussion

Reconstructing the demographic history of species from genetic data is a complex endeavor and a major challenge because many factors have likely influenced the genetic patterns we observe today. These factors include population structure, changes in connectivity and in population size, selection, social structure (mating systems), among others [11, 41, 24, 42]. Major progress in human population genetics and genomics, including paleogenomics, have revolutionized our understanding of present-day and past genetic variation [43, 44]. Ideas and methods coming from human population genetics have influenced our understanding of the genetic diversity of other species [45, 46, 6, 14]. Here we propose to rather use what we know about other great apes to ask questions about ancient population structure and the evolutionary history of humans, and perhaps learn from such comparative analyses, as others have also suggested [47]. The work presented here is thus both an attempt at increasing our understanding of the ancient structure of chimpanzees but also on the evolutionary history of humans.

### 4.1 Towards a demographic history of chimpanzees incorporating population structure

We managed to explain patterns of genetic diversity within each subspecies of common chimpanzees with simple models of population structure and variable migration rates only, and we obtained demographic scenarios for each subspecies with shared periods of connectivity change. The validation step we applied (see Supplementary Material) suggests that if real genomic data had been generated under the inferred demographic scenarios, our method would have been able to infer them. While we do take inferred scenarios with a grain of salt, this validation step suggests that complex population structure can be inferred from single genomes. This is particularly notable because we recovered the simulated scenarios for each subspecies independently. Though this does not confirm that chimpanzees evolved under the inferred scenarios, it suggests that SNIF can infer different complex scenarios from the chimpanzee PSMC curves. A similar validation process is often applied in ABC studies [39, 48] but it is not that common in the literature, and we argue that it should be implemented more often to validate scenarios proposed on the basis of other inference methods.

Despite some variability in the inferences, we observed consistency across inferences within and between subspecies. We found that Western chimpanzees are characterized by a higher number of demes with a smaller size than the other three subspecies. This could be consistent with the fact that this subspecies lives in a drier habitat, mostly in savanna [49], leading to a forest habitat that is more fragmented. However, it is unclear if it was the case throughout the evolutionary history of Western chimpanzees, and thus drawing strong conclusions is difficult at this stage. More generally, interpreting the number of islands and their size is not trivial and relating these parameters to empirical observations is not straightforward, in the same way that effective population size (*N_e_*) values inferred in previous studies assuming panmixia are not easily interpreted. We assumed for simplicity that the number of demes and their size were constant. Allowing for the number of demes to vary would be more realistic but would also significantly increase the number of parameters in the models. At this stage, the inferred values of *n* and *N* should thus be interpreted with care.

Our models suggest that the four subspecies share a similar history of connectivity until approximately 500 kya (forward in time). We found a period of high connectivity between 2.5 and 1 Mya, followed by a decrease in connectivity until approximately 600-500 kya. The start of this period coincides with the Pliocene-Pleistocene transition boundary dated to around 2.6 Mya, whereas the drop in connectivity around 1 Mya falls within the Middle Pleistocene transition thought to have occurred between 1.2 Mya and 700 kya [50]. A second and more recent period of high connectivity, between 600-500 and 200-150 kya is also observed, although in Western and Nigeria-Cameroon chimpanzees only. The fact that only two chimpanzee subspecies were affected by this increase in migration during the most recent Middle Pleistocene period suggests that whichever environmental disturbances caused the genetic signal observed in the PSMC, these disturbances were localised mostly in Western Africa. Interestingly, Mazet *et al.* [28] and Arredondo *et al.* [31] also inferred high values of migration rates between 2.5 and around 1 Mya using PSMCs from humans, but did not identify the more recent period of high connectivity found in Nigeria-Cameroon and Western chimpanzees. This suggests that humans may share a common environmental history with Eastern and Central African chimpanzees, rather than with Western and Nigeria Cameroon chimpanzees for this period, at least. This also supports the idea that comparative analyses of genomic data from other vertebrates and primates from Africa (Eastern and Western) could improve our understanding of the history of the genus *Homo* in the last two millions of years.

We must however be careful in interpreting genomic data and environmental changes together, and more work would be needed to validate these results since they are based on a simple interpretation of the PSMC curves and on the assumption that PSMC curves infer the IICR with sufficient precision. At this stage we still lack a clear demographic model that would integrate the Pliocene-Pleistocene and Middle Pleistocene transitions and that can explain how environmental changes would have affected habitat connectivity (or changes in effective population size), similarly impacting the PSMC curves in humans and chimpanzees. Also, we must acknowledge that reconstructing paleo-environments and habitats is still a complex endeavour [51, 52].

From a more technical point of view, we should also note that the inferred migration rates for Western chimpanzees were lower than for the other three species (Figure 4). This is surprising since all subspecies should provide similar values for the periods where there was one ancestral species to all four, and the PSMC curves overlap. We currently have no simple interpretation, and this could be due to some specificity of the Western subspecies, the quality of its genome, or the fact that we also inferred smaller deme sizes.

Similarly, we must note that the PSMC curves exhibit a large variance in the recent past with an apparent increase (forward in time) in the recent past that is interpreted in the connectivity graph as an increase in gene flow. We stress that this observed increase (forward in time) could also be due to a recent increase in the deme size, or to an uncertainty in the inference of the IICR. Indeed, the large variance in PSMC estimates in the recent past has been noted since the publication of the method of Li and Durbin [24]. Since SNIF assumes models with constant size it cannot typically fit this section of the IICR without making major changes in *M*. One must recall that under a model of constant size the IICR will necessarily stay at a low value (corresponding to the deme size in the recent past). IICR theory also shows that the IICR will “move up” quicker (backward in time) from the deme size to large values when *M* is large. Thus, under the assumption of constant size, SNIF will infer large *M* values in the very recent past to allow the IICR to move “up” rapidly from the inferred deme size, as explained by Mazet *et al.* [28]. At this stage, we thus considered that the recent increase in migration rate inferred for all chimpanzees should be interpreted very cautiously, if not ignored.

### 4.2 The general model and the limits of tree models

Using the demographic scenarios inferred for each subspecies of common chimpanzees, we successfully integrated the results of SNIF for each subspecies within a general tree model inspired by previous research on the four subspecies [2, 6, 14]. The difference with previous research was thus that we used *n*-island models instead of panmictic populations within each of the branches of the phylogenetic tree. We found that this model explained the PSMC curves of the four subspecies, and predicted well observed heterozygosity within demes and genetic differentiation between demes from the same subspecies. However, even using the shortest split times that would be consistent with the PSMC curves, the model led to an overestimation of genetic differentiation between subspecies. This suggests that the statistics that depended on the SNIF inference were generally better than those that depended on the tree model. This consequently implies that using a tree topology without gene flow to define the relationships between the four subspecies and ignoring possible gene flow between them may not be appropriate. Two recent studies identified IBD patterns across the four subspecies considered as a single unit, suggesting that the four subspecies were part of a very large spatial metapopulation with gene flow between neighbouring populations, including populations currently attributed to different subspecies. Spatial models, instead of *n*-island models, might thus be necessary to represent the evolutionary history of chimpanzees.

The idea that gene exchange may have taken place between subspecies has been present in the literature [2, 14, 53, 6, 13, 11, 12, 10]. However, in most cases gene flow was seen as discrete events that could be dated, or that were limited to a pair of subspecies, which were in some cases not in geographical contact. Brand *et al.* [14] reviewed the literature on this question and found that at least 14 admixture events had been identified by eight different studies (Figure 1 of Brand *et al.* [14]) including one admixture event from a mysterious ghost species into the ancestors of bonobos [53]. Among these putative admixture events, some were identified by only one study whereas others were identified by two to five. It also appeared that some studies identified only one admixture event [7] whereas others identified as many as eight [13]. Brand *et al.* [14] themselves used an inferential method (Legofit, [54]) and a model that allowed for up to seven admixture events but only found support for two.

Altogether, this suggests that it has been difficult to find a consistent history of admixture or gene flow among previous genetic studies. We must stress that these studies are not always easy to compare, and some differences may arise from the fact that their sampling was different. For instance, several studies have no sample from one or two chimpanzee subspecies, whereas in other cases, the authors used samples from individuals with unknown geographic origin. However, one common feature of all these studies is that they consider tree models that ignore population structure below the subspecies level. They usually assume that the chimpanzee subspecies and the bonobos should be modelled as independent and panmictic lineages of an evolutionary tree where the only gene flow allowed is through these discrete admixture events.

### 4.3 The limits of the piecewise *n*-island model: towards spatial models

The approach used throughout the manuscript follows the theoretical and simulation-based work of several authors who found that structured stationary and non-stationary models can generate genetic signatures that will be interpreted in terms of population size change when population structure is ignored [32, 55, 33, 56, 57, 28]. However, the *n*-island models we assumed with SNIF ignore spatial processes. As a consequence, IBD patterns observed by [17] and [18] cannot be reproduced, and suggest that the different subspecies were genetically connected in the recent past even if the chimpanzees’s habitat is currently highly fragmented and discontinuous.

A difficulty that could arise from the use of spatial models is that the parameter space may increase significantly making the inference process more difficult. However, if we wish to improve our understanding of the evolutionary history of great apes, including humans, we may have no choice but integrate spatial models [58, 59]. As a simple test example and to illustrate the importance of spatial models, we developed a simple 1D stepping-stone model inspired by the demographic model proposed by de Manuel *et al.* [6] to study the demographic history of chimpanzees and bonobos. These authors assumed a tree model and allowed for the possibility of admixture events between subspecies and between bonobos and chimpanzees. They computed D-statistics and found evidence for admixture between bonobos and chimpanzees. Details of our 1D stepping-stone models and most of the results can be found in the Supplementary Material, but here we mainly wish to stress that we were able to reproduce the D-statistics with our spatial model without any introgression between bonobos and chimpanzees. By changing gene flow and deme size in the structured ancestral species, we found that we were recovering even higher D values than those observed today. This suggests that the bonobo admixture signals detected in chimpanzees might be the simple result of both ancient and recent population structure.

## 5 Conclusions

In this work, we showed that it is possible to propose a demographic model for common chimpanzees that accounts for population structure and gives a coherent interpretation of PSMC curves produced by previous studies. Although we stress that the general model we propose here should not be taken as face value, it manages to explain several patterns of genetic diversity within subspecies despite the limits of the *n*-island models (e.g. lack of spatial attributes). We noted the importance of using spatial models to account for the genetic differentiation between the subspecies and also showed that spatial models might also explain possibly spurious signatures of admixture with bonobos. This work is a first step towards more complex models, though we recognise the difficulty of such endeavour. There is an increasing recognition that ignoring population structure and spatial processes may lead to the inference of events that may never have happened during the evolutionary history of the species studied [33, 36, 37, 60, 38]. This has implications for many species but also important consequences on the understanding of human evolutionary history.

## Ethics Statement

Not applicable.

## Consent for Publication

Not applicable.

## Availability of Data and Material

All code used for this paper is available at https://github.com/camillesteux/structurechimpanzees.

## Author Contributions

Conceptualization: LC; Methodology: LC, CS, CC, AA, WR, OM, RT; Investigation: LC, CS; Visualization: LC, CS, RT; Supervision: LC; Writing—original draft: LC, CS, RT ; Writing—review: CC, AA, WR, OM.

## Acknowledgments

We thank all the members of the Population and Conservation Genetic group at the Instituto Gulbenkian de Ciência (IGC) for their support and for insightful discussions on the topic. We also thank the Bioinformatics Unit and the Informatics Team of the IGC for their help and support with computational resources, and the Genotoul bioinformatics platform. Finally, we thank Mimi Arandjelovic, Jack Lester and Javier Prado-Martinez for sharing their data with us.

## Supplementary data

Supplementary Information S1: On the validation step.

Supplementary Information S2: Spatial structure confounds ancient admixture estimates.

Table S1: Prior ranges for the *t_i_*.

Table S2: Distribution of inferred *n*.

Table S3: Distribution of inferred *N*.

Figure S1: Model and statistics calculated under a no-admixture structured population model.

Figure S2: PSMC curves provided by Prado-Martinez et al..

Figures S3-S6: Inferred IICR curves for each subspecies of chimpanzees.

Figure S7: Distribution of inferred n and N coloured by individual.

Figure S8: Inferred connectivity coloured by individual.

Figure S9: Pseudo-observed data given to SNIF for the validation step.

Figure S10: Results of the validation step when giving to SNIF simulated PSMC curves as pseudo-observed data.

Figures S11-S14: Empirical PSMC and simulated IICR under the general tree model for each species of chimpanzees.

Figures S15-S17: Genetic distances within and between subspecies computed on genomic data simulated under the general tree model for different splitting times.

