## Supplementary Material for "On the demographic history of chimpanzees and some consequences of integrating population structure in chimpanzees and other great apes"

### Supplementary Information S1: On the validation process

The validation process that we applied in this study is similar to that used in ABC studies. We started with a PSMC curve for which we inferred ten scenarios corresponding to ten runs of SNIF, as explained in the Materials and Methods section. When these ten scenarios were similar we chose one of them, say scenario $S^*$, and generated a new IICR curve using the corresponding *ms* command. This IICR curve served as input to a new SNIF inference and we obtained ten newly inferred scenarios , $S_1^{**}, S_2^{**}, ...S_{10}^{**}$ from the ten independent runs. If these scenarios are different from each other, this may suggest that the optimization has not reached equilibrium and the number of iterations should be increased. Assuming now that the $S_1^{**}, S_2^{**}, ...S_{10}^{**}$ do not differ significantly from each other but differ from $S^*$, this suggests that the inferred scenario $S^*$ should not be trusted at this stage since SNIF was not able to infer it despite using ten independent runs and reaching equilibrium. This was never the case with the chimpanzee data and we come back to this below.

In such a case where $S^*$ should not be trusted, we suggest several solutions. One could explore more simple n-island models (with smaller $c$ values for instance) to determine if $S^*$ is too parameter-rich to be inferred by SNIF. If this fails too, it could also indicate that the real evolutionary history may be too complex to be approximated by a piecewise stationary n-island model such as $S^*$ or simpler versions of $S^*$. In such a case, other demographic models should be explored, perhaps involving population size changes or spatial structure (stepping stone models for instance), or tree models.

If, on the contrary, $S^*$ can be inferred reasonably well (i.e. the $S_i^{**}$ scenarios are similar to $S^*$) as we observe with the chimpanzees, this could suggest that the scenario $S^*$ is not only able to explain the original observed data but it can be inferred by SNIF if it were true. In other words, if the real species had evolved under $S^*$ we would be able to infer $S^*$ with SNIF. This does not prove that the species evolved under $S^*$ but that $S^*$ might be a reasonable approximation of reality to explain the PSMC computed for that species, at least until we find better alternatives.

These (or other) validation steps are fundamental and we suggest that they should be applied more often in demographic inference studies. It is more commonly applied in ABC studies, but there are still studies which infer a scenario without demonstrating that if real data had been generated under the inferred scenario, the authors would indeed be able to infer it back again, even approximately. While this validation process may be difficult to apply to some methods using genomic data, the current study shows that it is possible.

### Supplementary Information S2: Spatial structure confounds ancient admixture estimates

Several studies have inferred ancient admixture events between lineages of the *Pan* genus [6, 13, 11, 21] (cf. Brand *et al.* [14] for a review). The mode and tempo of these putative events vary greatly with little consistency across studies (see the Discussion section in the main manuscript). This could suggest a complex admixture history involving the different subspecies and the bonobos, of which previous research works identified only some elements. Alternatively, the inferred admixture events might also be caused by the confounding effect of population structure (or other departures from model assumptions) which could generate different results across studies, depending on the models, statistics, samples, etc. used by the authors.

For instance, de Manuel *et al.* [6] inferred introgression between chimpanzees and bonobos using different approaches including the *D*-statistic [61], `TreeMix` method, and *SFS*-based demographic inference (Site Frequency Spectrum). However, like most previous studies, they assumed panmixia within each subspecies, and thus did not test for ancient population structure, which is an increasingly recognised confounder for the detection of admixture [61, 36, 37, 38].

Here, we implemented a simple linear stepping-stone model to test the hypothesis that population structure alone, without any gene flow between bonobos and chimpanzees, could replicate the observed *D*-statistics. We found that non-zero *D*-statistics could indeed be generated, following a gradient similar to what is observed on the empirical data. Noteworthy, our model not only predicted the empirical levels of $D$ but it also predicted realistic values of the nucleotide diversity and the $F_{ST}$ among chimpanzee subspecies. Altogether, this suggests that population structure can reproduce several signals of present-day genetic diversity, including purported admixture ones. It thus calls for more caution regarding the strength of the evidence favoring ancient admixture over population structure in the extant *Pan* genus, in a way that is similar to that suggested in recent research on the genus *Homo* [36, 37, 38].

#### S2.1 Methods

##### S2.1.1 Demographic model

We implemented a simple linear (one-dimensional) stepping-stone model (Fig. S1A), where each subspecies of chimpanzee is treated as a "metapopulation" of five connected demes of respective size $N_i$ diploids. Bonobos are modelled in a similar way, except that no bonobo subspecies is currently recognised. We will abbreviate henceforth: Western chimpanzees as "W", Nigeria-Cameroon as "NC", Central as "C", Eastern as "E", and Bonobos as "P" (for *paniscus*). Thus, the full model is composed of five metapopulations of five demes each.

For simplicity, we set the deme size in each metapopulation using the $\theta_W$ estimate produced by de Manuel *et al.* [6] (Table S2 in the original article), with $N = \theta_W/(4n\mu)$, where $n$ is the number of demes we consider in this study for each metapopulation ($n = 5$). We used the same mutation rate as de Manuel *et al.*, *i.e.* $\mu = 1.2 \times 10^{-8}$ per bp per generation. This led us to set $N_{NC} = 5{,}559$ diploids, $N_E = 6{,}498$, $N_C = 9{,}462$. For Western chimpanzees, we found that the original $\theta$ (leading to $N_W = 3{,}475$) produced an excess of nucleotide diversity compared to empirical values and we thus set it to a lower value ($N_W = 600$). We fixed the deme size in bonobos to $N_P = 4{,}000$ (about half $N_C$ [62]), in the absence of the $\theta_W$ estimate in de Manuel *et al.* [6].

Within all metapopulations, demes are connected to their neighbours with a per-generation symmetric migration rate of $m_w = 10^{-2}$ (*i.e.* $M_{i,j} = m_w N_i$ diploid individuals migrating from deme $i$ to deme $j$ at each generation, backward). The different chimpanzee metapopulations are connected to the neighbouring metapopulation by a low-level (symmetric) gene flow of $m_b = 8 \times 10^{-4}$. The migration rate within the bonobo metapopulation was fixed at $m_{w,P} = 1.1 \times 10^{-4}$.

We acknowledge that several of these parameter values were fixed arbitrarily, since our purpose was not to infer parameters. We intended to showcase how a simple model can explain empirical data (when accounting for known population structure, without requiring unknown ancient admixture events), especially in producing apparent signatures of putative ancient admixture events.

Metapopulations split times were implemented in the model using the average of the divergence times reported in de Manuel *et al.* [6] (Figure 3 in the original article). Specifically, NC started to expand from W at 250 kya; E from C at 160.5 kya, the two ancestral chimpanzee lineages at 588.5 kya, bonobos from chimpanzees at 1.88 Mya. Each expansion is modelled, forward in time, by successive founding of each within-metapopulation deme, every 200 years. We note that these founding events correspond exclusively to the instantaneous creation of a deme of size $N_i$ (*i.e.* no bottlenecks).

For the time period prior to the first split between chimpanzee metapopulations (588.5 kya), the deme sizes in the ancestral chimpanzees were set to $N_{anc} = 7,000$ and the migration rate between the demes of this ancestral metapopulation was set to the same value as the present-day bonobo migration rate (*i.e.* $m_{anc} = m_{w,P} = 1.1 \times 10^{-4}$). All parameters for the remaining bonobo metapopulation were kept the same.

It should be clear that this model assumes that bonobos and chimpanzees never exchange gene flow at any time of their history after they separated 1.88 Mya.

The total number of non-redundant parameters in our model is 15, making it less parameter-rich than the panmictic model of de Manuel *et al.* [6]:

- 1 parameter for the number of demes per metapopulation (set to 5),
- 2 parameters for within-metapopulation migration rate in chimpanzees and bonobos (set to $m_w = 10^{-2}$ and $1.1 \times 10^{-4}$, respectively),
- 1 parameter for between-metapopulation migration rate in chimpanzees (set to $m_b = 8 \times 10^{-4}$),
- 5 parameters for deme sizes ($N_i$),
- 1 parameter for ancestral deme size ($N_{anc}$),
- 4 expansion times (*i.e.* the founding of the different metapopulations),
- 1 parameter for the delay between successive founding of demes (set to 200 years).

##### S2.1.2 Simulations

Using `msprime v1.1` with Hudson's coalescent algorithm [63, 64], we simulated genetic data for 20 chromosomes of 20 Mbp each ($G = 400$ Mbp), sampling 15 diploid individuals in the central deme of each metapopulation. This resulted in a total of 75 diploid genotypes per segregating site. Mutations were generated using a binary model, with two flipping alleles, at a rate of $1.2 \times 10^{-8}$ per bp per generation. Recombination was assumed uniform, with rate $0.7 \times 10^{-8}$ per bp per generation. We used a generation time of 25 years for both chimpanzees and bonobos [6].

##### S2.1.3 Statistics

Based on the simulated biallelic genetic data, we calculated, using the `scikit-allel v1.3.5` package:

- The nucleotide diversity $\pi$ for each population sample (average number of pairwise differences).
- The differentiation index $F_{ST}$ (Hudson's formula based on expected heterozygosities) between all pairs of chimpanzee populations.
- The *D*-statistic, as $D(X,Y;bonobo,outgroup)$ with $X$ and $Y$ being any chimpanzee population sample, and *outgroup* being a virtual diploid genotype homozygous for the ancestral allele. To calculate the standard error (*SE*), we used a block-weighted jackknife with a typical block size of around 5 Mbp [65]. Confidence intervals at 95% were calculated as $D \pm 1.96 \times SE$.

##### S2.1.4 Observed data

We retrieved the empirical statistical values from previously published papers:

- $\pi$ (average pairwise differences): from Fischer *et al.* [22], calculated on 26 intergenic sequences totalling 22.4 kbp for around 10 diploids in each sampled population.
- $F_{ST}$ (Hudson's formula): from Fischer *et al.* [22], calculated on the same dataset as $\pi$.
- *D*-statistic: from de Manuel *et al.* [6], calculated as $D(X,Y,bonobo,humans)$ using whole-genome sequences on a set of 68 *Pan* samples. We used the *D* values estimated by the authors when aligning the *Pan* sequences on the human *hg19* assembly.

#### S2.2 Results

Our 1D stepping-stone structured population model replicated the observed gradient of *D*-statistics across focal chimpanzee subspecies (Fig. S1D). It further predicted the empirical values of between-chimpanzee $F_{ST}$ (with the Central-Eastern being slightly underestimated in our simulations) (Fig. S1C), as well as the nucleotide diversity, with a slight excess of diversity for Eastern chimpanzees (Fig. S1A) (which likely results from the $N_E$ value that we extracted from [6]).

In conclusion, these results show that the ancient admixture inferred in the *Pan* genus might not be robust to ancestral population structure.

We further investigated the variation of the $D$-statistic as a function of the ancestral connectivity, i.e. the connectivity within the metapopulation ancestral to extant chimpanzees (and further, extant *Pan* species) from 588.5 kya towards the past. To this end, we sampled $N_{anc}m_{anc}$ values from $10^{-3}$ to 10, every 0.2 on a $\log_{10}$-scale. Keeping $N_{anc}$ fixed at 7,000 (cf. previous model), we derived $m_{anc}$ according to the composite parameter $N_{anc}m_{anc}$. All other demographic and simulation parameters were kept the same as in the previous model, except for genome size, restricted to $10 \times 10$ Mbp chromosomes for computational reasons. We estimated $D$ between present-day Central and Western chimpanzees: $D(C, W, bonobo, outgroup)$.

Our results (Fig. S2) show that the $D$-statistic values follow a sigmoid curve, "saturating" at zero for the highest $N_{anc}m_{anc}$ values (towards the right). This is expected, since with such high connectivity, the population model becomes nearly panmictic, and it does not incorporate admixture from bonobos into C lineages. The curve also plateaus around 0.6 for the lowest $N_{anc}m_{anc}$ range. Interestingly, we note that the $D$-statistic values start to become significant, in our model, around $N_{anc}m_{anc} = 1$ migrant per generation. A steep variation in $D$ is observed for connectivity values ranging between around 0.06 and 0.6 migrants per generation. These results confirm that very significant values of the $D$-statistic can be produced under a structured population model in the absence of admixture, and that they depend mostly on the level of the ancestral connectivity within the metapopulation ancestral to the tested samples. We show that $D$ can reach very high values (0.6) compared to the empirical values reported here, confirming that the level and significance of the $D$ statistic cannot be used as indisputable evidence for admixture.

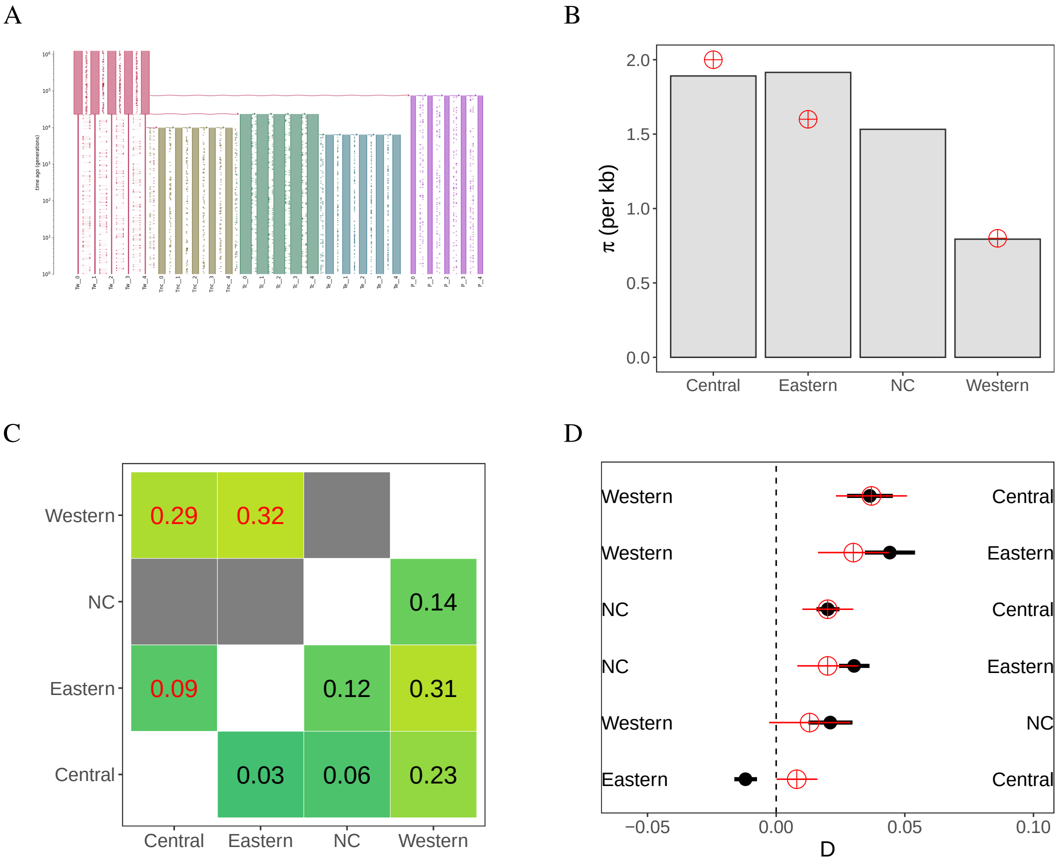


Figure S1: Model and statistics calculated under a no-admixture structured population model. A. Representation of the simulated demographic model. Each color corresponds to a chimpanzee subspecies or to the bonobos. Red: W, olive: NC, green: C, blue: E, purple: bonobos. B. Nucleotide diversity per kbp estimated for each population, for the data simulated under the structured population model (gray bars), and on the empirical data (red points). No NC samples were available in the [22] study. C. Pairwise $F_{ST}$ obtained on the simulated data (lower-right triangle, black text) and on the empirical data (upper-left triangle, red text). We note that in the original article, the NC population was not studied [22], thus appearing here as empty gray cells. D. $D$-statistics from the simulated data (black points with 95% confidence interval) and empirical data (red points). The $D$-statistic values were calculated as $D(X,Y,P,O)$ with "P" the bonobos. On the plot, $X$ populations are labeled on the right and $Y$ on the left.

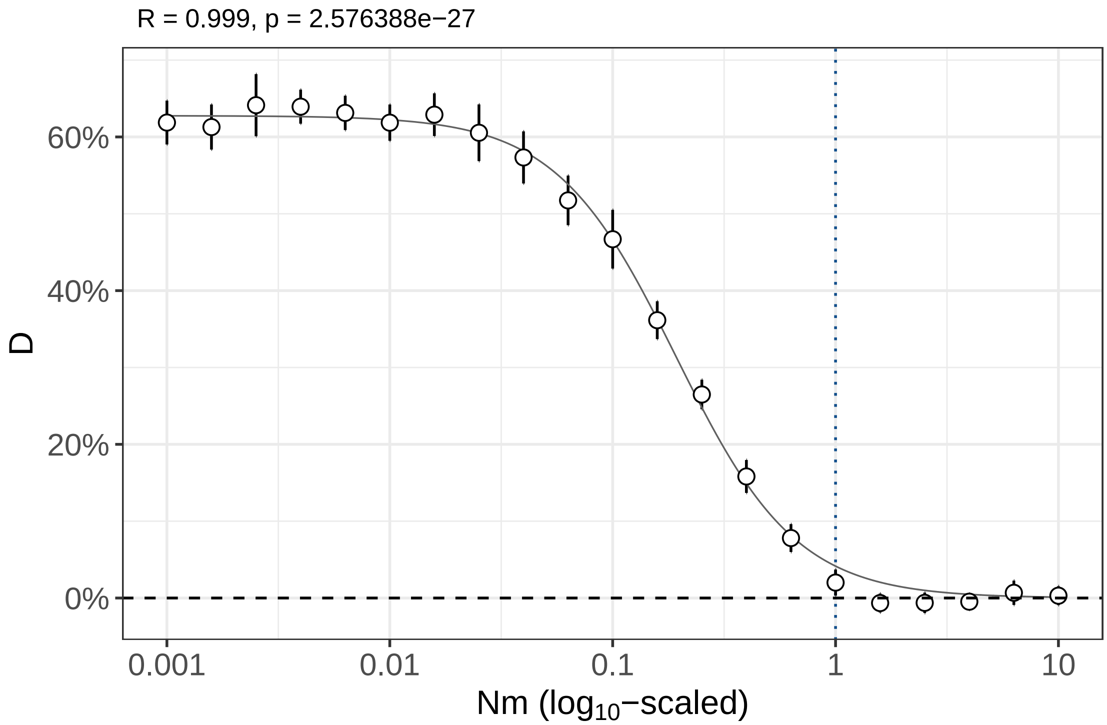


Figure S2: Distribution of the $D$-statistic as a function of ancestral migration rates. The $D$-statistic was computed as $D(C,W,P,O)$ for our no-admixture structured model, where we allowed the migration rate in the ancestral metapopulation to vary. This ancestral metapopulation corresponds to the period prior to 588.5 kya, *i.e.* before the foundation of the bonobos (cf. model description) We allowed $N_{anc} \times m_{anc}$ to vary from $10^{-3}$ to 10. The error bars represent the confidence intervals at 95%. The $x$-axis is $\log_{10}$-scaled. The vertical blue dotted line represents the first tested $N_{anc}m_{anc}$ value from decreasing order with significant $D$-statistics. The gray curve is a logistic fit to the empirical scatter plot. The Pearson's correlation coefficient representing the fit of the curve to the empirical data is reported in the title.

Table S1: Prior ranges of the $t_i$ (times at which migrations rates are allowed to change, delimiting the components) given to SNIF for the final analysis. Note that here the times are given in years, but they have to be divided by the generation time when given to SNIF.

| Subspecies | $c$ | Priors of $t_i$ (in years) |
|---|---|---|
| Western | 7 | (2e4, 1e5), (1e5, 3e5), (1e5, 3e5), (3e5, 7e5), (7e5, 1.5e6), (1.5e6, 5e6) |
| Nigeria-Cam. | 7 | (1e4, 1e5), (1.5e5, 3e5), (3.5e5, 5e5), (8e5, 1.1e6), (2e6, 3e6), (4e6, 7e6) |
| Central | 8 | (5e4, 2e5), (2e5, 4e5), (5e5, 2e6), (5e5, 2e6), (2e6, 5e6), (2e6, 5e6), (6e6, 1e7) |
| Eastern | 7 | (5e4, 2e5), (4e5, 1.5e6), (4e5, 1.5e6), (2e6, 6e6), (2e6, 6e6), (7e6, 1e7) |

Table S2: Distribution of inferred $n$, the number of demes of the $n$-island models.

| **Subspecies** | **Min** | **25% quantile** | **Median** | **Mean** | **75% quantile** | **Max** |
|---|---|---|---|---|---|---|
| Western | 12 | 17 | 21 | 23 | 31 | 48 |
| Nigeria-Cameroon | 6 | 10 | 11 | 11 | 13 | 20 |
| Central | 13 | 16 | 18 | 21 | 20 | 55 |
| Eastern | 8 | 11 | 13 | 13 | 15 | 23 |

Table S3: Distribution of inferred $N$, the deme size of the $n$-island models (in number of diploids).

| **Subspecies** | **Min** | **25% quantile** | **Median** | **Mean** | **75% quantile** | **Max** |
|---|---|---|---|---|---|---|
| Western | 112 | 239 | 305 | 285 | 335 | 437 |
| Nigeria-Cameroon | 616 | 1009 | 1154 | 1174 | 1311 | 1980 |
| Central | 227 | 589 | 737 | 694 | 834 | 1101 |
| Eastern | 561 | 728 | 801 | 863 | 975 | 1306 |

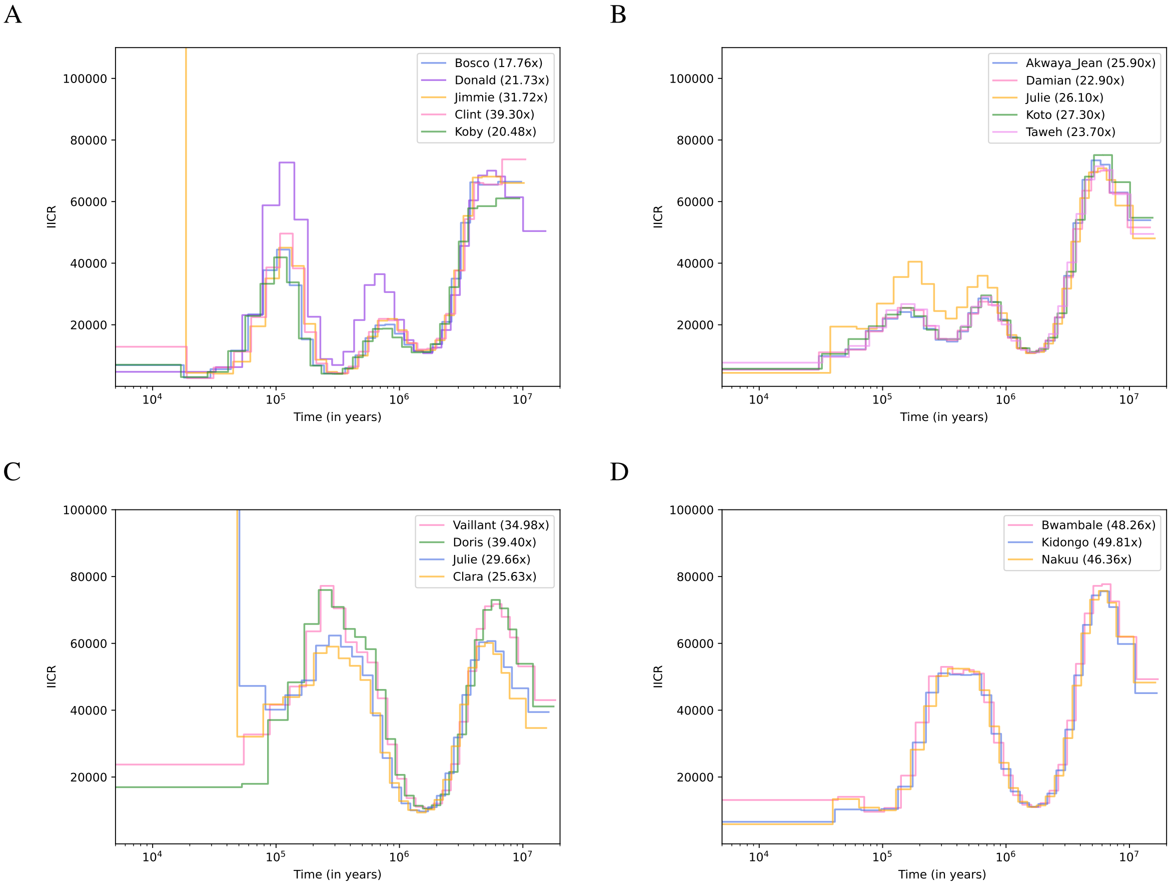


Figure S3: PSMC curves for A. Western chimpanzees, B. Nigeria-Cameroon chimpanzees, C. Central chimpanzees and D. Eastern chimpanzees, computed using PSMC files provided to us by Prado-Martinez *et al.* [2]

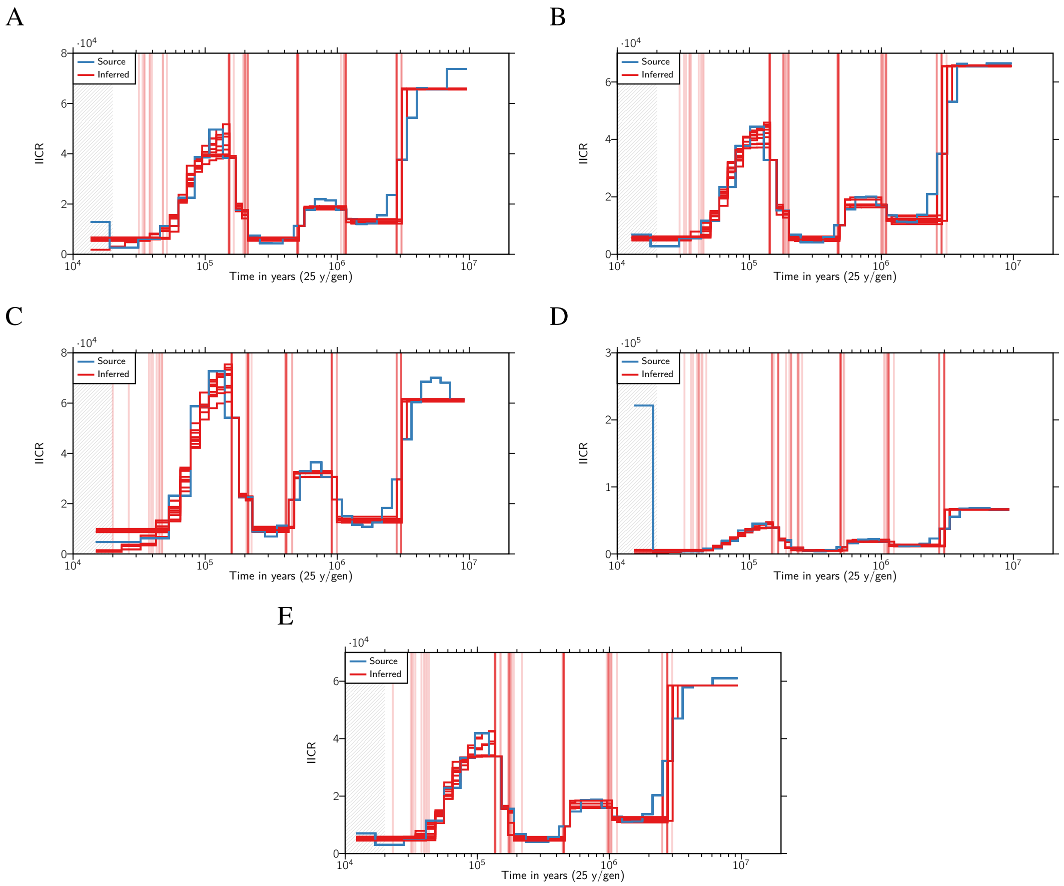


Figure S4: Inferred IICR curves (in red) and empirical PSMC (in blue) for the five Western common chimpanzees. Each red curve is a repetition of SNIF. The vertical red lines highlight the times at which there is an inferred change in migration rate and therefore delimit the components. A. Clint, B. Bosco, C. Donald, D. Jimmie and E. Koby. The grey zone corresponds to a part of the source PSMC which was not taken into account for the fitting of the curve by SNIF (see Material and Methods).

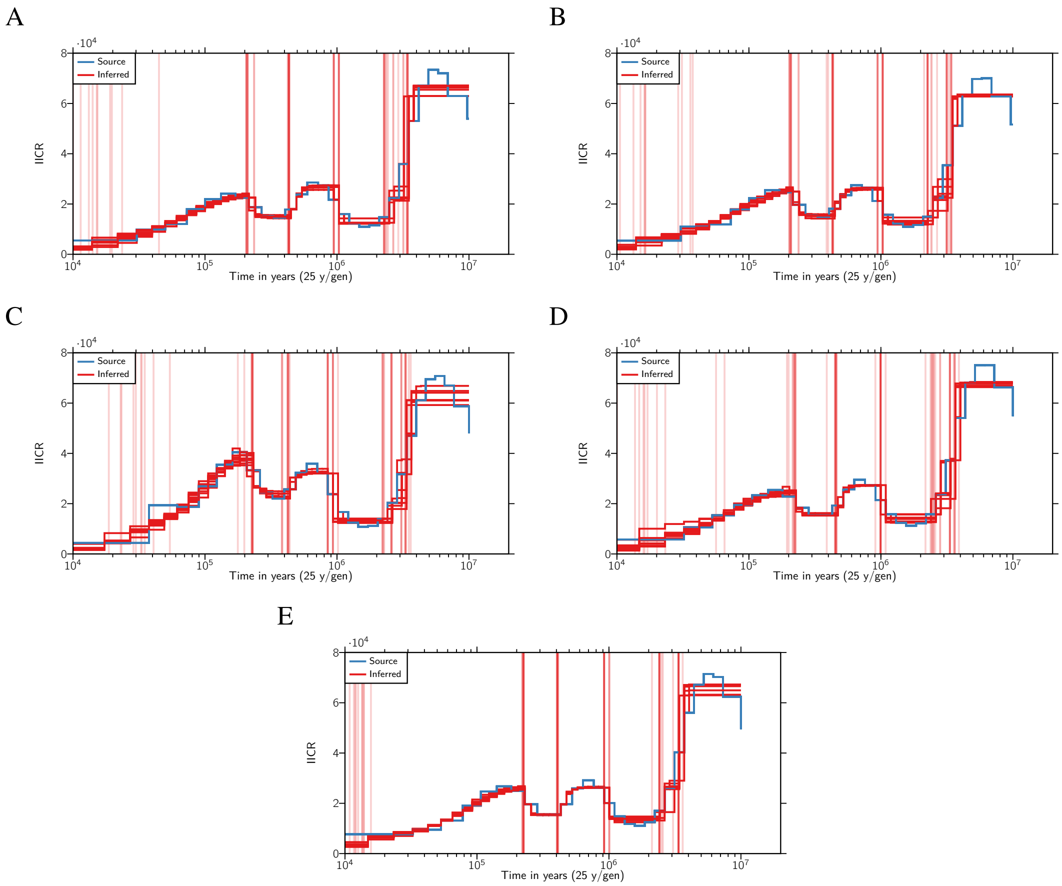


Figure S5: Inferred IICR curves (in red) and empirical PSMC (in blue) for the five Nigeria-Cameroon common chimpanzees. Each red curve is a repetition of SNIF. The vertical red lines highlight the times at which there is an inferred change in migration rate and therefore delimit the components. A. Akwaya-Jean, B. Damian, C. Julie, D. Koto and E. Taweh.

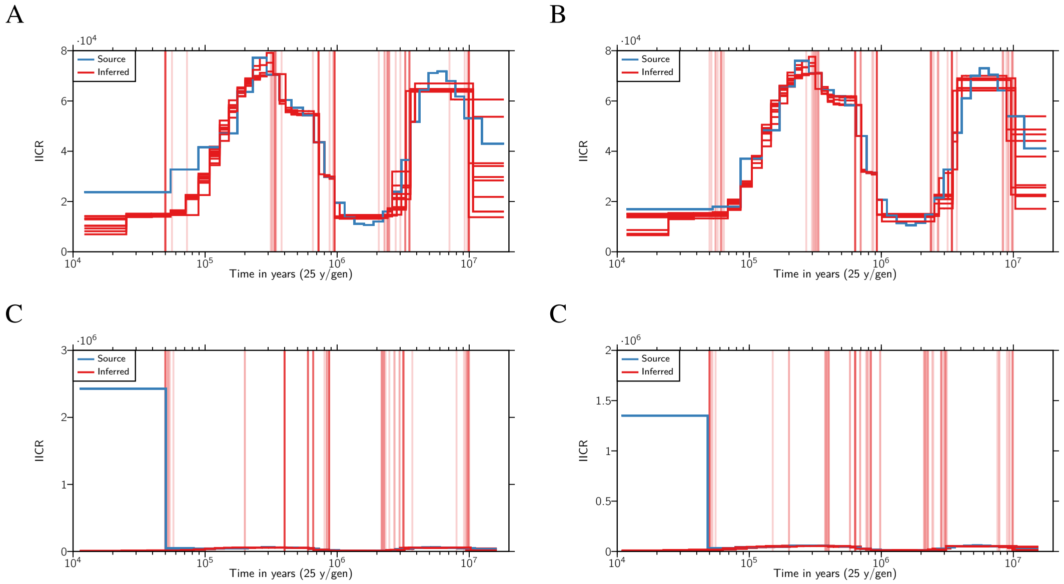


Figure S6: Inferred IICR curves (in red) and empirical PSMC (in blue) for the four Central common chimpanzees. Each red curve is a repetition of SNIF. The vertical red lines highlight the times at which there is an inferred change in migration rate and therefore delimit the components. A. Vaillant, B. Doris, C. Julie and D. Clara.

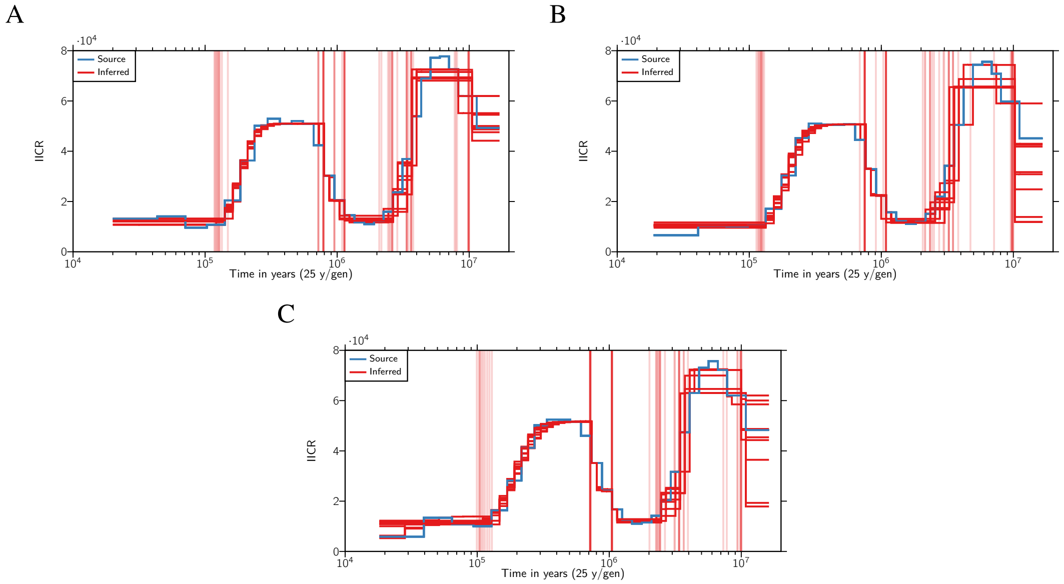


Figure S7: Inferred IICR curves (in red) and empirical PSMC (in blue) for the three Eastern common chimpanzees. Each red curve is a repetition of SNIF. The vertical red lines highlight the times at which there is an inferred change in migration rate and therefore delimit the components. A. Bwambale, B. Kidongo and C. Nakuu.

A

B

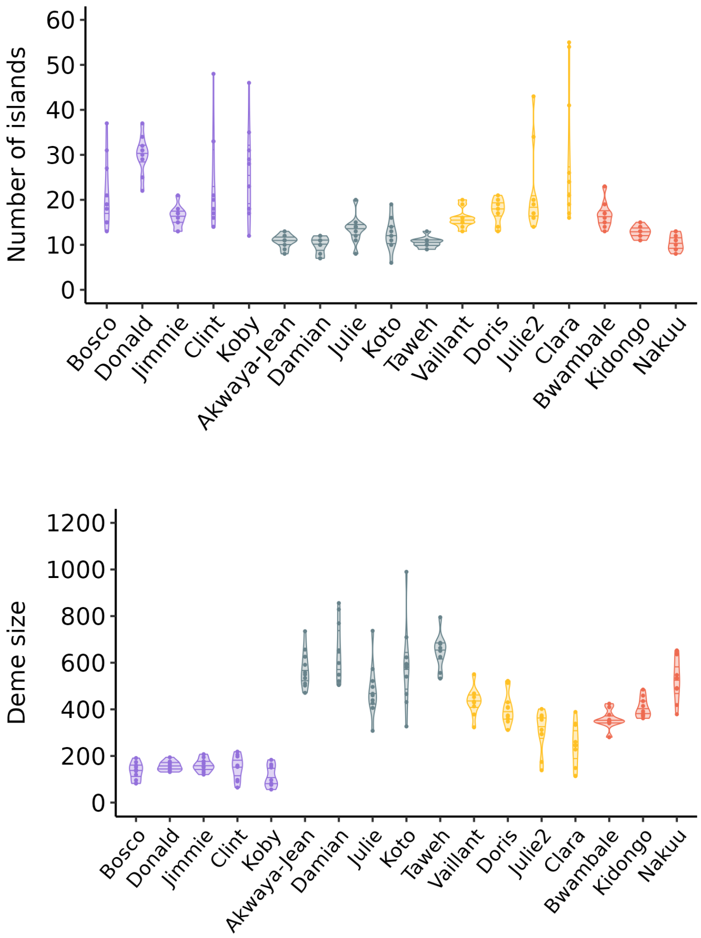


Figure S8: Distribution of the inferred A. number of islands and B. deme size (given in number of diploid individuals) across the 10 repetitions for each individual, using the parameter space shown Table 1. Horizontal lines in the violins represent the 25%, 50% (median) and 75% quantiles.

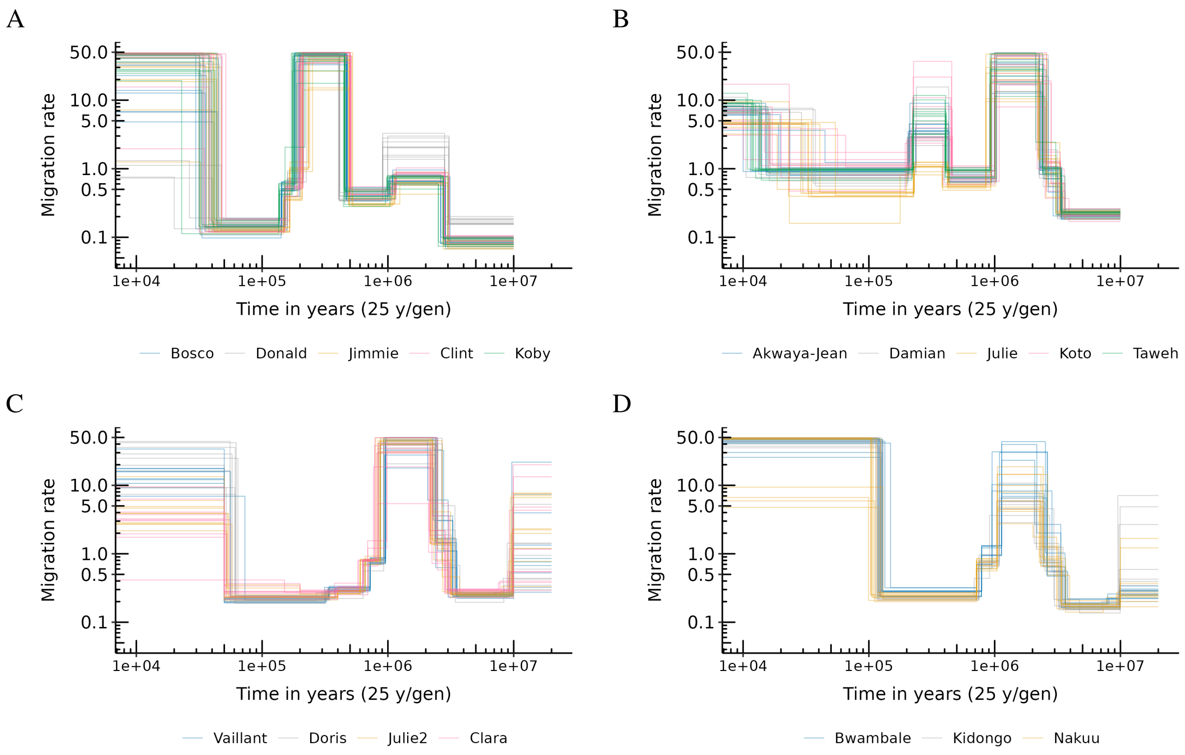


Figure S9: Connectivity graph (migration rates along successive time components) inferred by SNIF coloured by individual for A. Western, B. Nigeria-Cameroon, C. Central and D. Eastern chimpanzees. Each line corresponds to one inference (one repetition of SNIF) using one PSMC curve (or individual) as observed data.

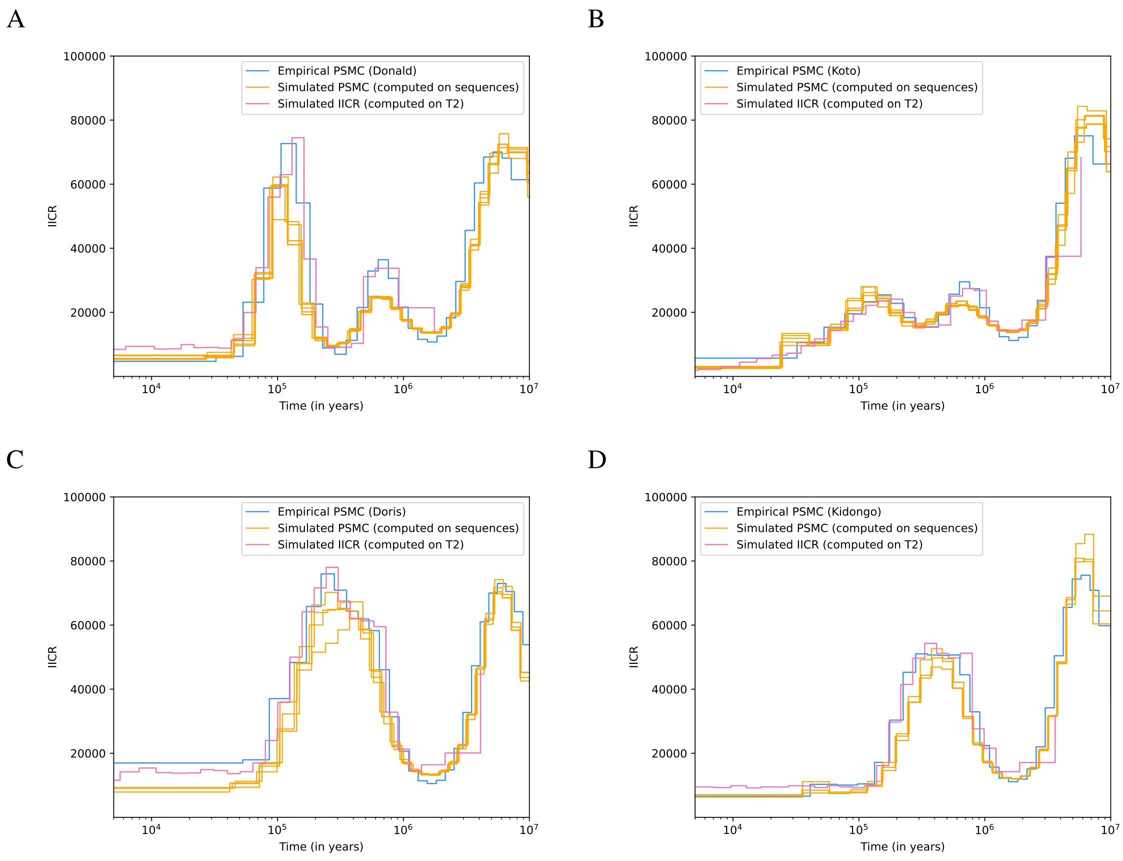


Figure S10: Simulated PSMC computed on simulated sequences (10x100Mb) (in blue) and simulated IICR computed on simulated coalescent times ($T_2$) (in orange), both given to SNIF as pseudo-observed data for the validation procedure. A. Western chimpanzees, B. Nigeria-Cameroon chimpanzees, C. Central chimpanzees and D. Eastern chimpanzees.

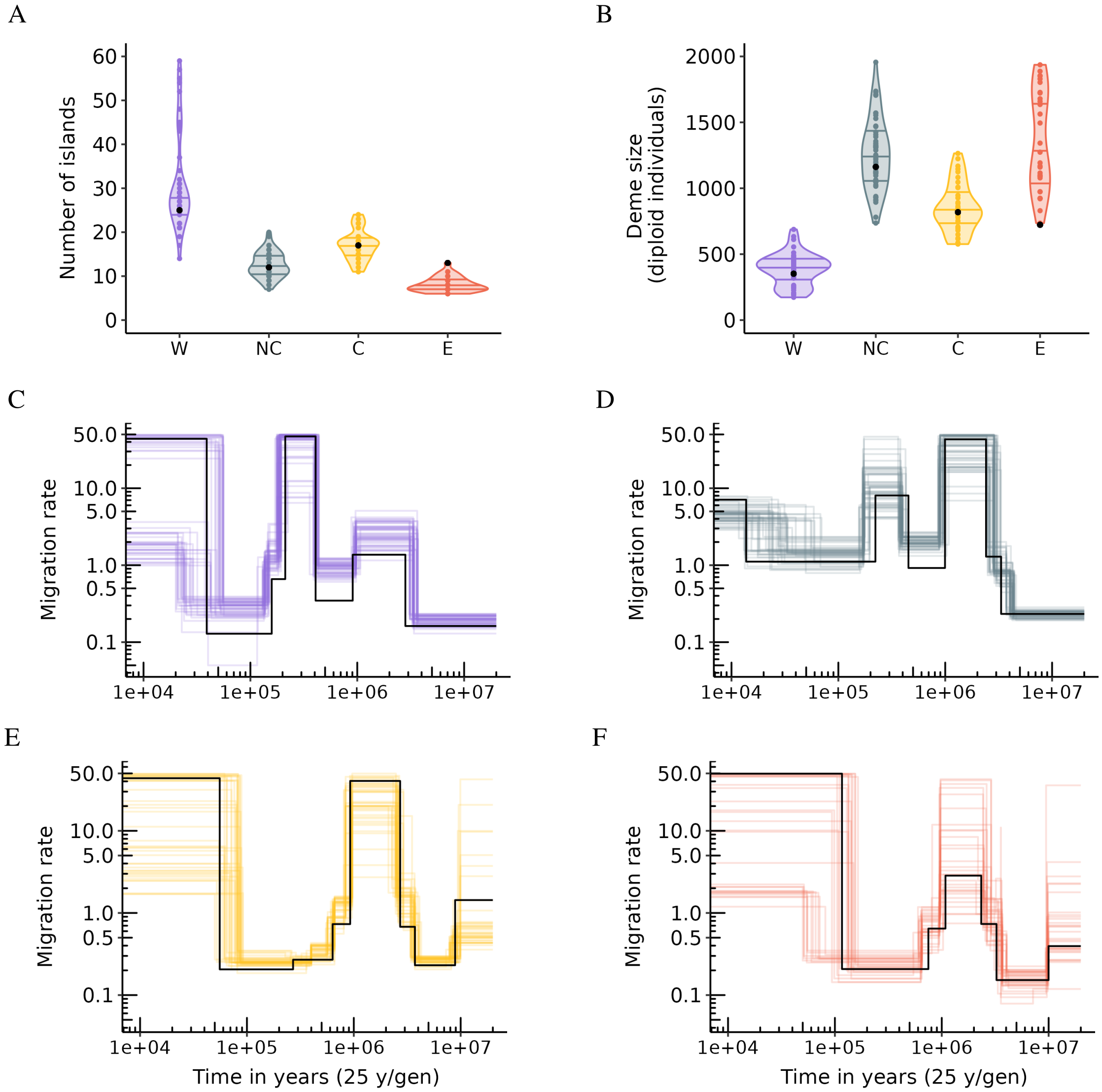


Figure S11: Results of the validation procedure when giving to SNIF simulated PSMC (see orange curves in Figure S10) as pseudo-observed data. A. Inferred number of islands ($n$), B. Inferred deme sizes ($N$), C. Inferred connectivity graph for Western chimpanzees, D. Inferred connectivity graph for Nigeria-Cameroon chimpanzees, E. Inferred connectivity graph for Central chimpanzees and F. Inferred connectivity graph for Eastern chimpanzees. Horizontal lines in the violins on panels A and B represent the 25%, 50% (median) and 75% quantiles.

A

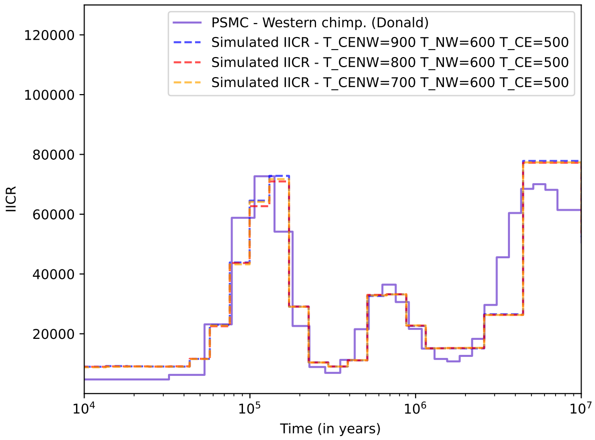


B

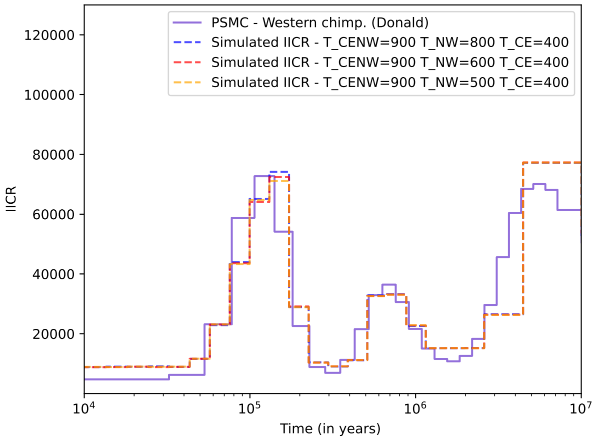


Figure S12: Empirical PSMC (solid line) and simulated IICR (dotted lines) for Western common chimpanzees under the general n-island model Figure 7 for different values of splitting times. (A) $T_{CENW} \in \{700, 800, 900\}$, $T_{NW} = 600$ and $T_{CE} = 500$ and (B) $T_{CENW} = 900$, $T_{NW} \in \{500, 600, 800\}$ and $T_{CE} = 400$ (in kya).

A

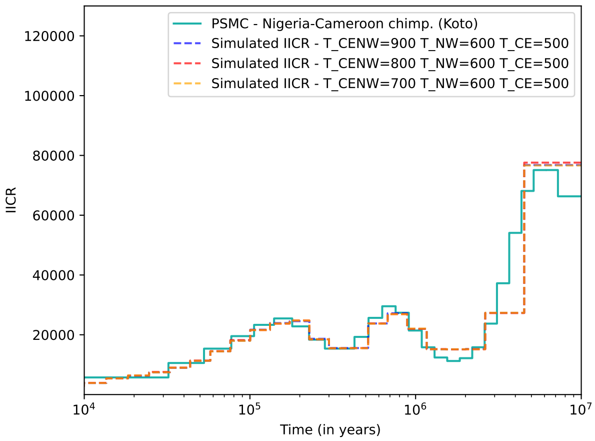


B

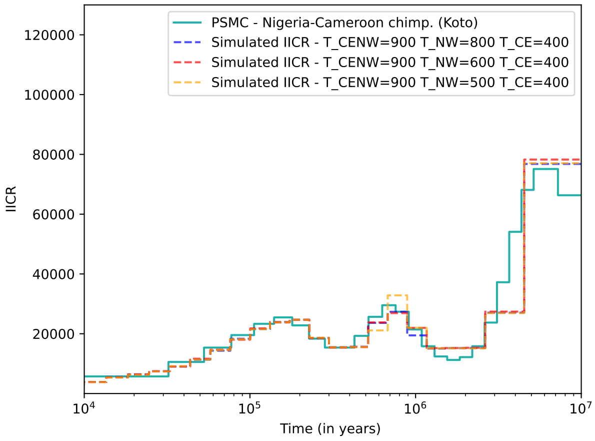


Figure S13: Empirical PSMC (solid line) and simulated IICR (dotted lines) for Nigeria-Cameroon common chimpanzees under the general n-island model Figure 7 for different values of splitting times. (A) $T_{CENW} \in \{700, 800, 900\}$, $T_{NW} = 600$ and $T_{CE} = 500$ and (B) $T_{CENW} = 900$, $T_{NW} \in \{500, 600, 800\}$ and $T_{CE} = 400$ (in kya)

A

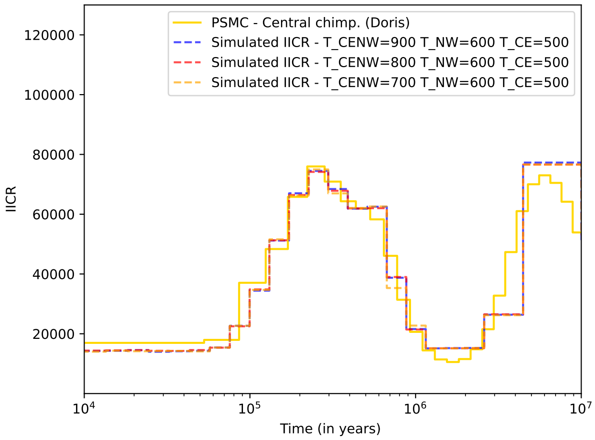


B

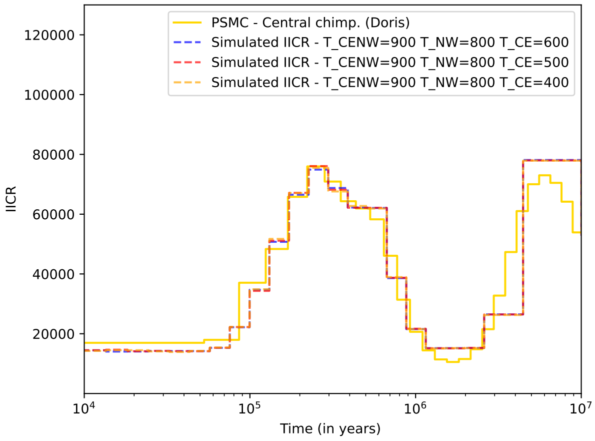


Figure S14: Empirical PSMC (solid line) and simulated IICR (dotted lines) for Central common chimpanzees under the general n-island model Figure 7 for different values of splitting times. (A) $T_{CENW} \in \{700, 800, 900\}$, $T_{NW} = 600$ and $T_{CE} = 500$ and (B) $T_{CENW} = 900$, $T_{NW} = 800$ and $T_{CE} \in \{400, 500, 600\}$ (in kya).

A

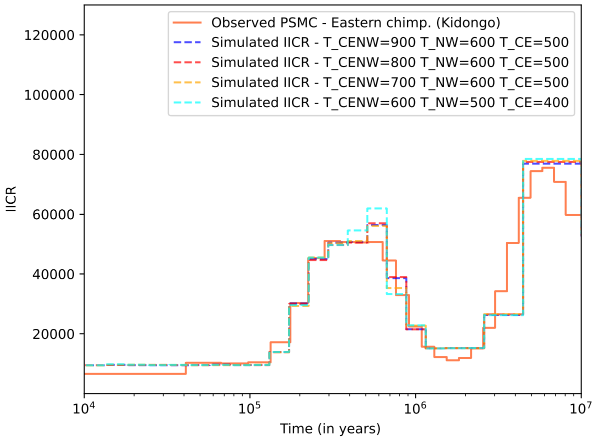


B

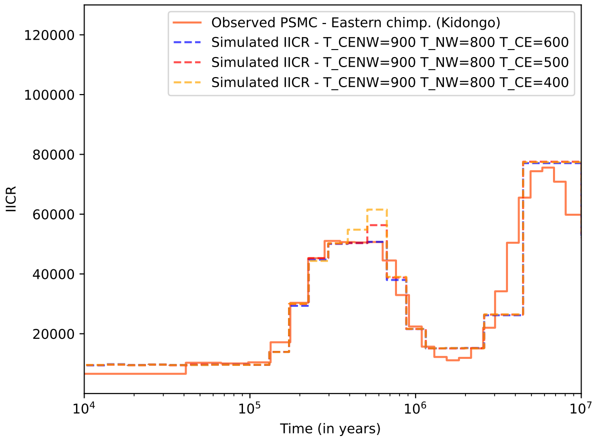


Figure S15: Empirical PSMC (solid line) and simulated IICR (dotted lines) for Eastern common chimpanzees under the general n-island model Figure 7 for different values of splitting times. (A) $T_{CENW} \in \{700, 800, 900\}$, $T_{NW} = 600$ and $T_{CE} = 500$ and (B) $T_{CENW} = 900$, $T_{NW} = 800$ and $T_{CE} \in \{400, 500, 600\}$ (in kya).

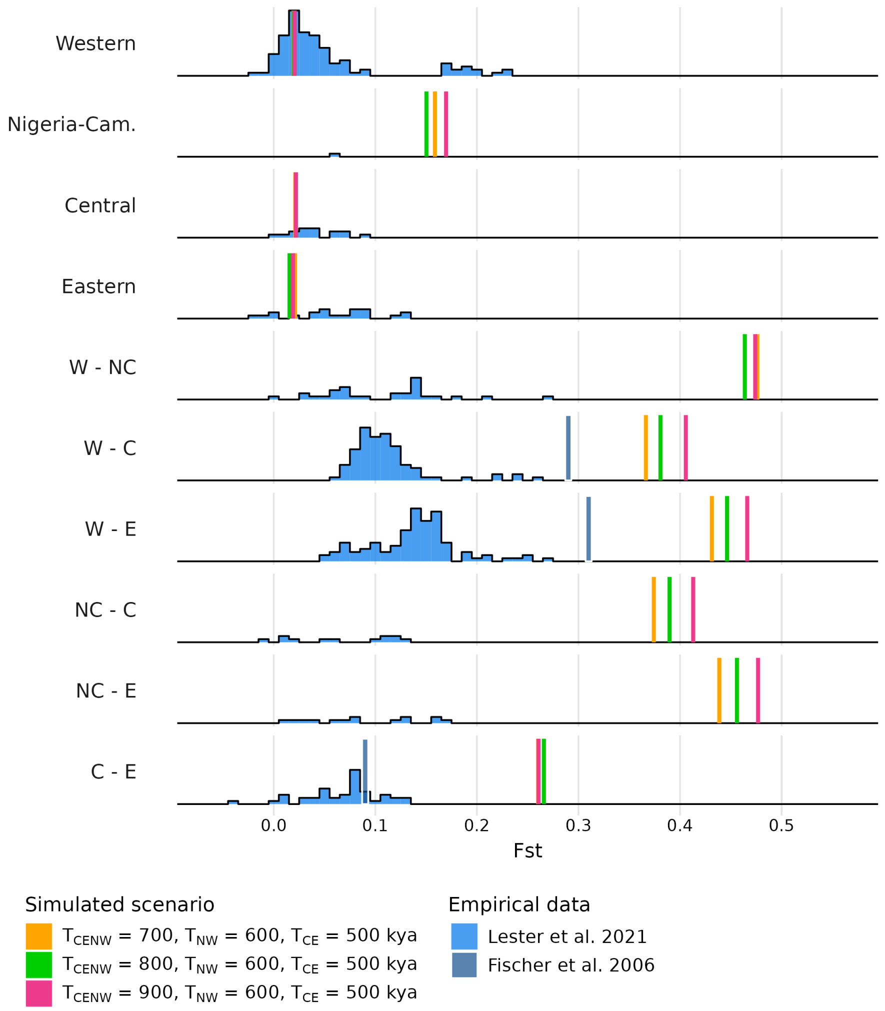


Figure S16: Genetic distance ($F_{ST}$) between demes of the same subspecies or between demes from different subspecies computed on genomic data simulated under the model Figure 7 with $T_{CE} = 500$ kya, $T_{NW} = 800$ kya and $T_{CENW} \in \{700, 800, 900\}$ kya. In blue are the empirical values: histograms were retrieved from Lester *et al.* [18] and the darker blue vertical lines were retrieved from Fischer *et al.* [22]

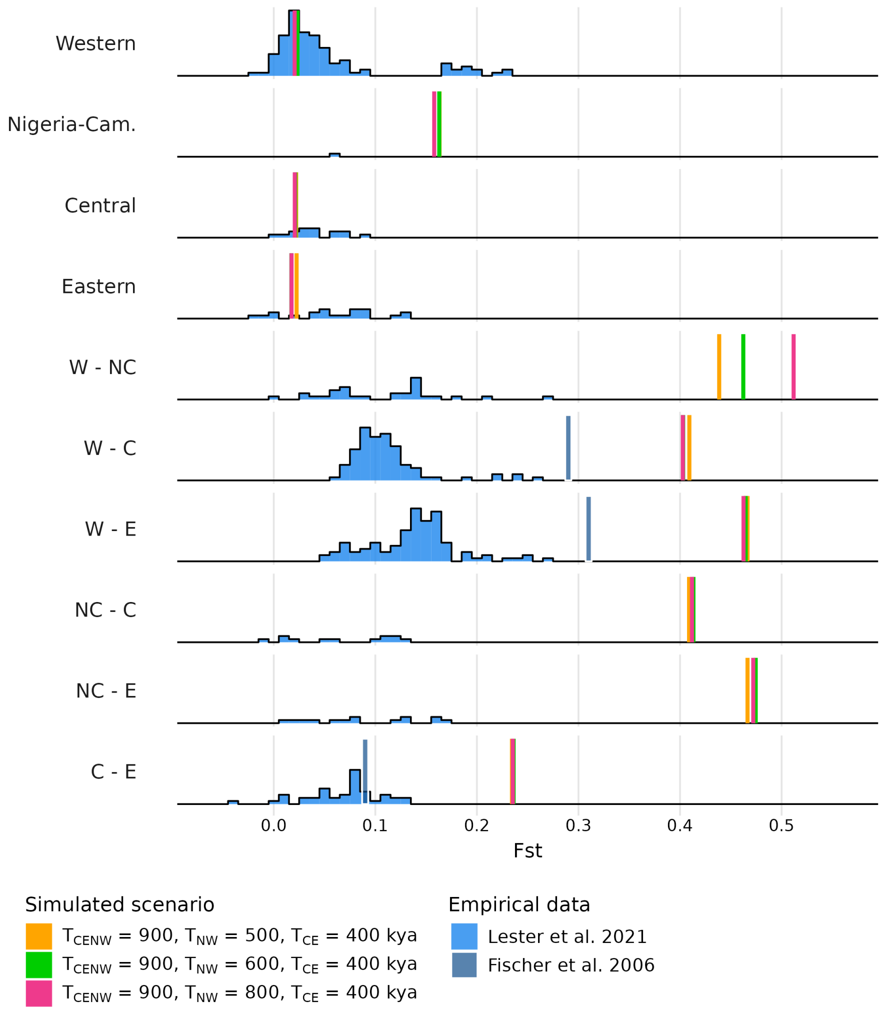


Figure S17: Genetic distance ($F_{ST}$) between demes of the same subspecies or between demes from different subspecies computed on genomic data simulated under the model Figure 7 with $T_{CE}$ = 500 kya, $T_{CENW}$ = 900 kya and $T_{NW} \in \{500, 600, 800\}$ kya. In blue are the empirical values: histograms were retrieved from Lester *et al.* [18] and the blue vertical lines were retrieved from Fischer *et al.* [22]

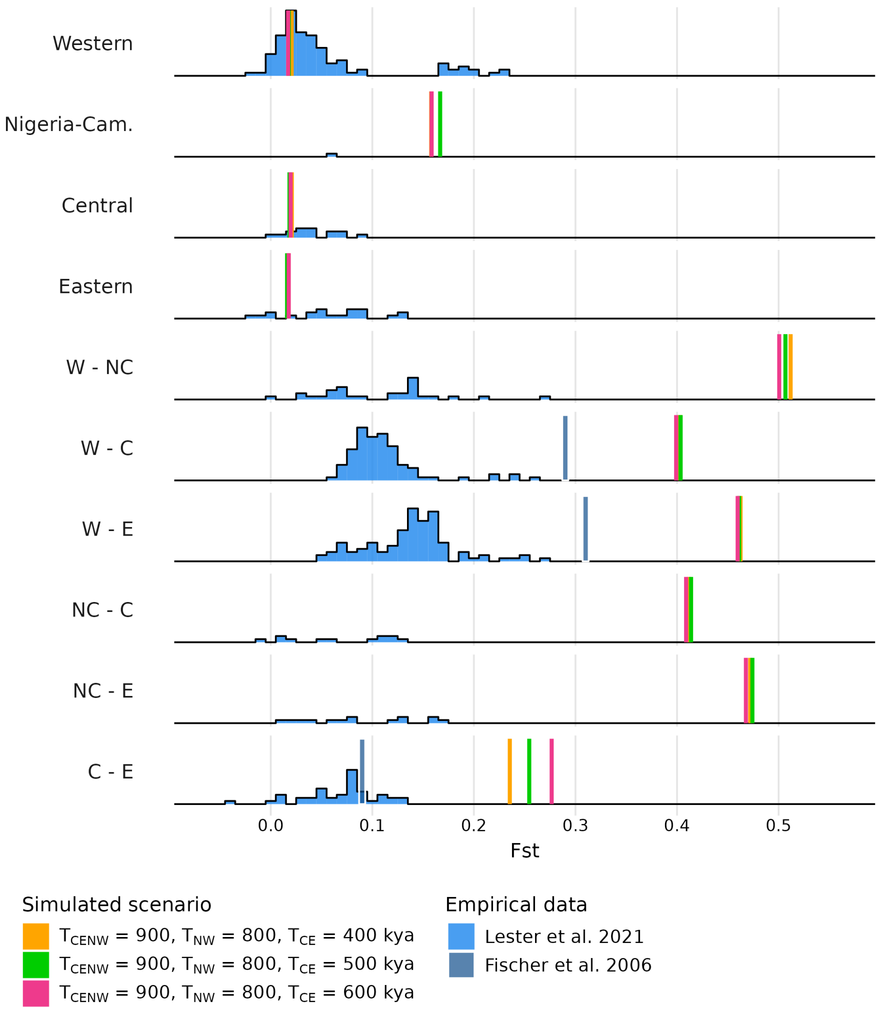


Figure S18: Genetic distance ($F_{ST}$) within and between subspecies computed on genomic data simulated under the model Figure 7 with $T_{CENW}$ = 900 kya, $T_{NW}$ = 800 kya and $T_{CE} \in \{400, 500, 600\}$ kya. In blue are the empirical values: histograms were retrieved from Lester *et al.* [18] and the blue vertical lines were retrieved from Fischer *et al.* [22]
